# *Cis*-epistasis at the *LPA* locus and risk of coronary artery disease

**DOI:** 10.1101/518290

**Authors:** Lingyao Zeng, Nazanin Mirza-Schreiber, Claudia Lamina, Stefan Coassin, Christopher P. Nelson, Oscar Franzén, Marcus E. Kleber, Salome Mack, Till F. M. Andlauer, Beibei Jiang, Barbara Stiller, Ling Li, Christina Willenborg, Matthias Munz, Thorsten Kessler, Adnan Kastrati, Karl-Ludwig Laugwitz, Jeanette Erdmann, Susanne Moebus, Markus M. Nöthen, Annette Peters, Konstantin Strauch, Martina Müller-Nurasyid, Christian Gieger, Thomas Meitinger, Elisabeth Steinhagen-Thiessen, Winfried März, Johan L. M. Björkegren, Nilesh J. Samani, Florian Kronenberg, Bertram Müller-Myhsok, Heribert Schunkert

## Abstract

Identification of epistasis affecting complex human traits has been challenging. Focusing on known coronary artery disease (CAD) risk loci, we explore pairwise statistical interactions between 8,068 SNPs from ten CAD genome-wide association studies (n=30,180). We discovered rs1800769 and rs9458001 in the vicinity of the *LPA* locus to interact in modulating CAD risk (P=1.75×10^−13^). Specific genotypes (e.g., rs1800769 CT) displayed either significantly decreased or increased risk for CAD in the context of genotypes of the respective other SNP (e.g., rs9458001 GG *vs*. AA). In the UK Biobank (n=450,112) significant interaction of this SNP pair was replicated for CAD (P=3.09×10^−22^), and was also found for aortic valve stenosis (P=6.95×10^−7^) and peripheral arterial disease (P=2.32×10^−4^). Identical interaction patterns affected circulating lipoprotein(a) (n=5,953; P=8.7×10^−32^) and hepatic apolipoprotein(a) (apo(a)) expression (n=522, P=2.6×10^−11^). We further interrogated potential biological implications of the variants and propose a mechanism explaining epistasis that ultimately may translate to substantial cardiovascular risks.

## Main text

Globally, coronary artery disease (CAD) is the largest contributor to morbidity and mortality^1^. Genetic understanding of CAD has benefited from recent genome-wide association studies (GWAS), which have identified multiple variants to independently and additively propagate CAD susceptibility^2^. Less attention has been paid to epistasis in which variants act non-additively or, paradoxically, are dependent on the genetic context^3,4^. Albeit epistasis has broadly been shown to affect multiple traits in fruit flies^5^, mice^6^, and humans^7^, examples demonstrating biological relevance for epistasis via statistical approaches have only been investigated in model organisms^8–10^. Earlier attempts were largely unsuccessful given the computational challenges such as the curse of dimensionality, model complexity and bias from linkage disequilibrium (LD). Moreover, such efforts often lacked successful replication and/or biological interpretations of the statistical interactions^11–15^. Here, we aimed to statistically identify pairwise SNP interactions in the context of CAD and explored how biological epistasis may functionally contribute to disease susceptibility.

We started from the search space of 56 broad-sense CAD susceptibility regions defined as ±500kb flanking regions of known CAD risk loci reported from previous CAD GWAS^16^ (Methods). Such focus was encouraged by the observation that respective regions contain substantial heritability not explained by respective lead SNPs (Supplementary Note I). Individual-level and imputed genotypes were utilized from 30,180 participants of ten European CAD case-controls studies^2,16–23^ (Methods, Supplementary Table 1, Supplementary Note II). We tested statistical interactions for all pairwise SNPs (n_SNP_LDpruned_=8,068) along a two-step scheme as detailed in the Methods and in Supplementary Fig 1. As a result, four SNP-pairs (Methods, Supplementary Table 2, Supplementary Fig 2), displaying consistent (in at least eight of ten studies) and significant (P ≤ 4.618×10^−9^) effects met our statistical criteria for candidate epistasis (Methods, Supplementary Fig 3, 4). The top SNP-pair (rs1800769- rs9458001) in a dosage-dosage model (Fig 1a, Supplementary Table 3) was prioritized for further investigation (Methods).

**Fig 1.**
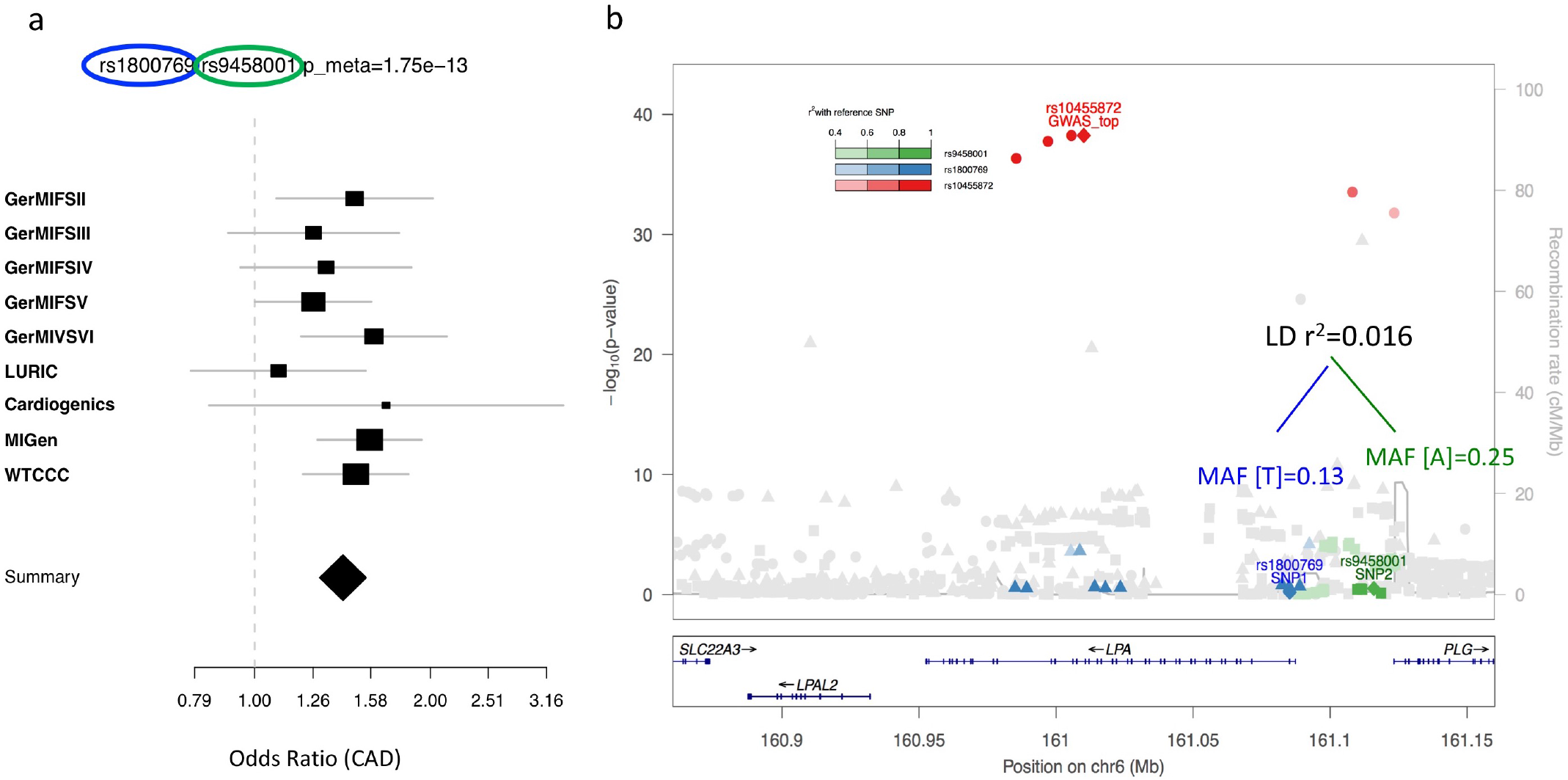
Cis-epistasis SNP-pair at the *LPA* locus displaying a non-additive signal increasing CAD risk. **a.** Forest plot displaying the effect sizes across 10 studies of the interaction term on CAD odds ratio between rs1800769[T] and rs9458001[A] as well as the meta-analysis summary effect (shown as a diamond). SNP rs1800769 was not available in the GerMIFSI study data, therefore it is not displayed in the forest plot. For this study, the summary statistics for the OR_int_ displayed the same trend, and the meta-analysis OR_int_ with perfect proxy SNPs across 10 studies is shown in (Supplementary Fig 4), with a fixed effect meta-analysis p-value as low as 1.75e^−13^. **b.** Manhattan plot displays the regional signals at the *LPA* locus taken from recent genome-wide association studies of CAD, with SNP rs1800769 (univariately p=0.08, OR=1.03, Supplementary Table 2) and those with a LD r^2^ > 0.4 are labelled in blue, SNP rs9458001 (univariately p=0.59, OR=0.99, Supplementary Table 2) and those with a LD r^2^ > 0.4 are labelled in green, and SNP rs10455872 (which was the so-far reported the top risk variant for CAD) and those with a LD r^2^ > 0.4 are labelled in red. LD: linkage disequilibrium. MAF: minor allele frequency. OR: odds ratio.

Both rs1800769 and rs9458001 map to chromosome 6, close to the apo(a) or *LPA* locus with a LD of *r*^2^=0.016 (Fig 1b). Neither were dosage-wise associated with CAD risk by itself (P=0.59, odds ratio (OR)=0.99 for rs1800769[T]; P=0.08, OR=1.04 for rs9458001[A]). However, together they displayed strong association (OR_int_=1.42, P=1.75×10^−13^ for the rs1800769[T]-rs9458001[A] interaction term) (Supplementary Table 2, Fig 1a, 1b). Statistical interaction was reliably present when conditioning for any known GWAS susceptibility SNPs for CAD (n=164) or any available SNP in the flanking ±200kb region. The same applies to known GWAS susceptibility SNPs for lipoprotein(a) (Lp(a)) (n=3)^2,24^ (Methods, Supplementary Table 4, 5, Supplementary Fig 5).

To facilitate comprehension, we investigated the relative odds ratio for subgroups of individuals with all nine possible genotype combinations. As a result, opposing genetic effects were observed for rs1800769, in that increasing dosage of T-alleles ([C][C]->[C][T]->[T][T]) went along with lower CAD risk in rs9458001[G] homozygote individuals, but with higher risk in rs9458001[A] homozygotes (Fig 2a). Comparable effects were observed but *vice versa* for rs9458001 given the different genetic context of rs1800769 (Fig 2a). The few double homozygote [T/T] - [A/A] genotype carriers were all found amongst CAD cases (Supplementary Note III). We further dissected the genetic context into four possible allele-specific subgroups of haplotype samples (Methods, Supplementary Fig 6). The allele combination [T-A] (∼3% in the population) displayed an odds ratio for CAD (1.84; Fig 2b), which was higher than that of any other common risk alleles at the Lp(a) locus, including rs10455872 at the Lp(a) locus^2,16^ (Supplementary Fig 7). Again, paradoxical effects were observed, where rs1800769[T] compared to [C] displayed lower risk in co-presence of rs9458001[G], but in contrast, displayed higher risk in co-presence of rs9458001[A] (Fig 2b, Supplementary Table 6). These strengthened the plausibility that the significant statistical interaction detected was due to the mutual dependency of genetic context (alleles) of two SNPs on the risk of CAD, and that there is true epistasis underlying the detected interaction signal.

**Fig 2.**
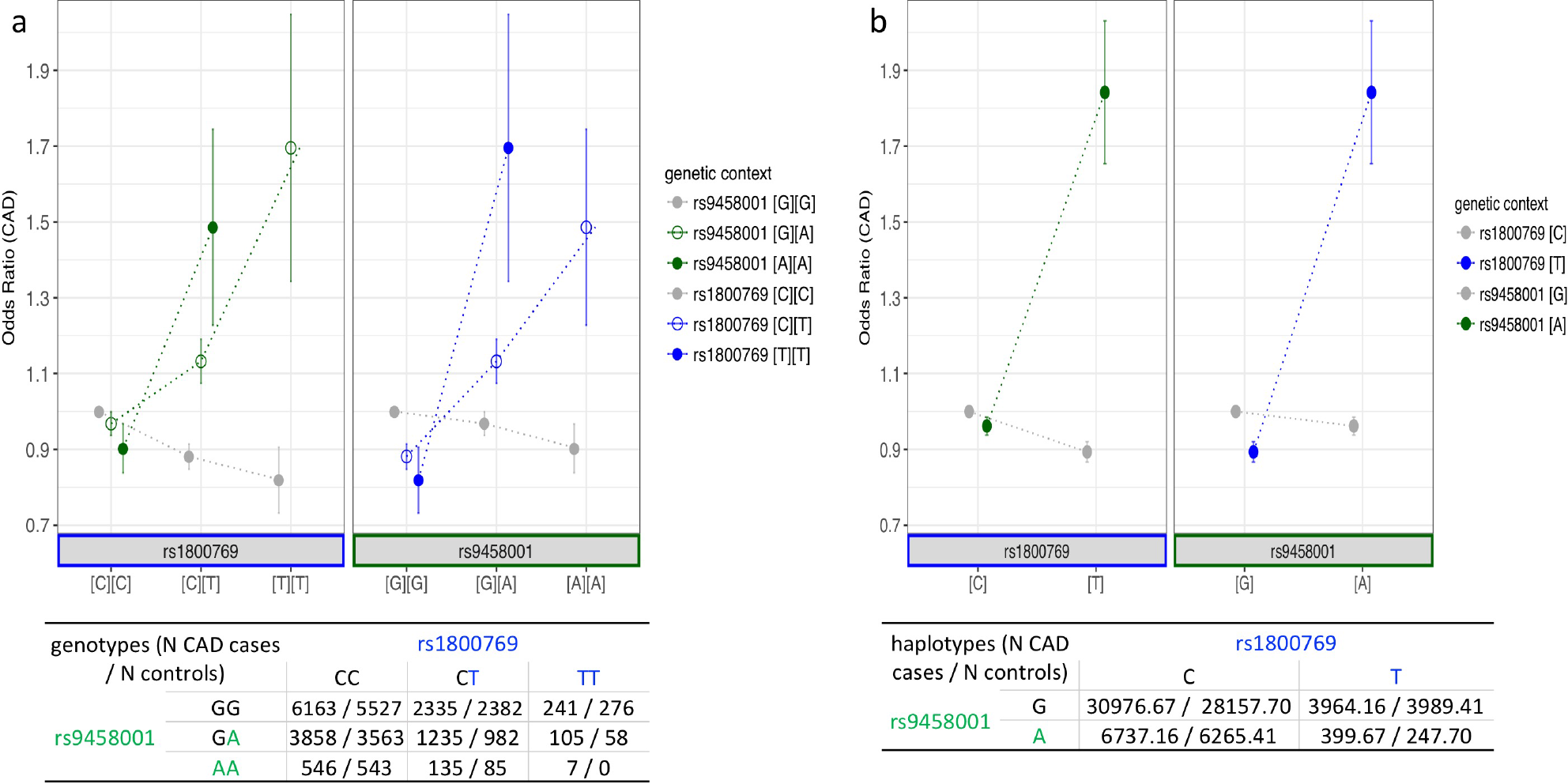
Statistical interaction between SNPs on CAD risk. **a.** Odds ratio of CAD and standard errors are displayed in biallelic perspective for 8 genotype combinations, with the reference genotype combination set as rs1800769[C][C]-rs9458001[G][G], the most frequent genotype. **Left panel:** With increasing numbers of [T] allele, ([C][C]➔[C][T]➔ [T][T], labelled in blue along x-axis) rs1800769 was related to lower CAD risk against the genetic context of rs9458001[G] homozygotes (along with the grey dotted line as viewing assistance). By contrast, CAD risk largely increased against the context of rs9458001[A] homozygotes (along with the green dotted line and solid points as viewing assistance), the effect was in-between against the genetic context of rs9458001 heterozygotes (along with the green dotted line and hollow points as viewing assistance). **Right panel:** Vice versa, with increasing numbers of the [A] allele, ([G][G]➔[G][A]➔[A][A], labelled in green along x-axis) rs9458001 was related to lower, largely increasing, and (in-between) mildly increasing CAD risk against the context of rs1800769[C][C], [T][T], and [C][T], respectively (lines and dots colored in grey or blue as viewing assistance). Bottom panel: Numbers of individuals were given corresponding to the subgroups of genotype combinations. **b.** Relative odds ratio of CAD and standard errors are displayed for 4 haplotypes, with the reference haplotype set as rs1800769[C]-rs9458001[G]. First pseudo samples of all possible haplotypes were generated compatible with each individual’s genotype, with appropriate weighting where haplotypes were uncertain. In addition haplotype association analyses were conducted based on generalized linear models using the expectation-maximization (EM) algorithm (Methods). **Left panel:** The haplotypes are shown in two groups based on the rs1800769[T] allele (marked in blue along the x-axis). The odds ratio of CAD was slightly reduced against the genetic context of rs9458001[G] (along with the grey dotted line and points as viewing assistance), but on the opposite was drastically increased against the genetic context of rs9458001[A] (along with the green dotted line and points as viewing assistance). **Right panel:** Vice versa, in the presence of rs9458001[A] odds ratios of CAD were either slightly reduced or drastically increased against the context of rs9458001[G] and [A], respectively. Bottom panel: Numbers of pseudo samples were given corresponding to the subgroups of haplotypes (allele combinations).

We replicated the statistical interaction of the two SNPs for CAD in two independent cohorts (UK Biobank n=443,588, P=3.09×10^−22^; GerMIFSVII n=5,379, P=6.7×10^−3^; meta-analysis across all studies with OR_rs1800769_=0.99, OR_rs9458001_=1.05, while OR_int_=1.30, P=5.07×10^−27^, Supplementary Table 7). Furthermore, in the UK Biobank data we found interaction effects in the same direction and comparable magnitude on peripheral arterial disease (controls/cases n=475,059/4,460, OR_rs1800769_=0.94, OR_rs9458001_=1.03, while OR_int_=1.22, P=2.32×10^−4^) and aortic valve stenosis (controls/cases n=477,496/2,023, OR_rs1800769_=0.94, OR_rs9458001_=1.00, while OR_int_=1.47, P=6.95×10^−7^, Supplementary Table 7), both of which are other manifestations of atherosclerosis in coronary arteries in which Lp(a) plasma levels affect risk^25,26^.

We next studied potential intermediary traits. Given that Lp(a), a well-known risk factor for CAD, is under strict genetic control (exceeding 90% in the European population)^27–29^, and is directly encoded at the locus near to rs1800769 and rs9458001, we analyzed circulating Lp(a) levels in a German population-based study (KORA F3/F4^24,30^, n=5,953). Despite a relatively trivial association with each SNP separately, we identified a strong interaction effect of both SNPs on Lp(a) level (beta=0.58, P=8.7×10^−32^) (Supplementary Table 6). Again, given the context of one SNP, paradoxical effects were observed for the other SNP (Fig 3, Supplementary Fig 8). Up to 3.0% of the variance of serum Lp(a) was explained by different genotype subgroups. Directional effects on Lp(a) levels and CAD risk were highly correlated for the genotype combinations (*r*= 0.96, P=3×10^−4^, Methods). Moreover, in the LURIC study where 2,831 individuals had both data on CAD onset and multiple lipid and coagulation markers, we replicated significant statistical interaction for Lp(a) levels, but found no other circulating factor displaying such effect (data not shown). In this cohort, the interaction coefficient on CAD risk was attenuated by adjustment for Lp(a) levels (Supplementary Table 8). These data plausibly suggest that the SNP-pair affects Lp(a) serum concentration, which subsequently results in higher risk of atherosclerosis.

**Fig 3.**
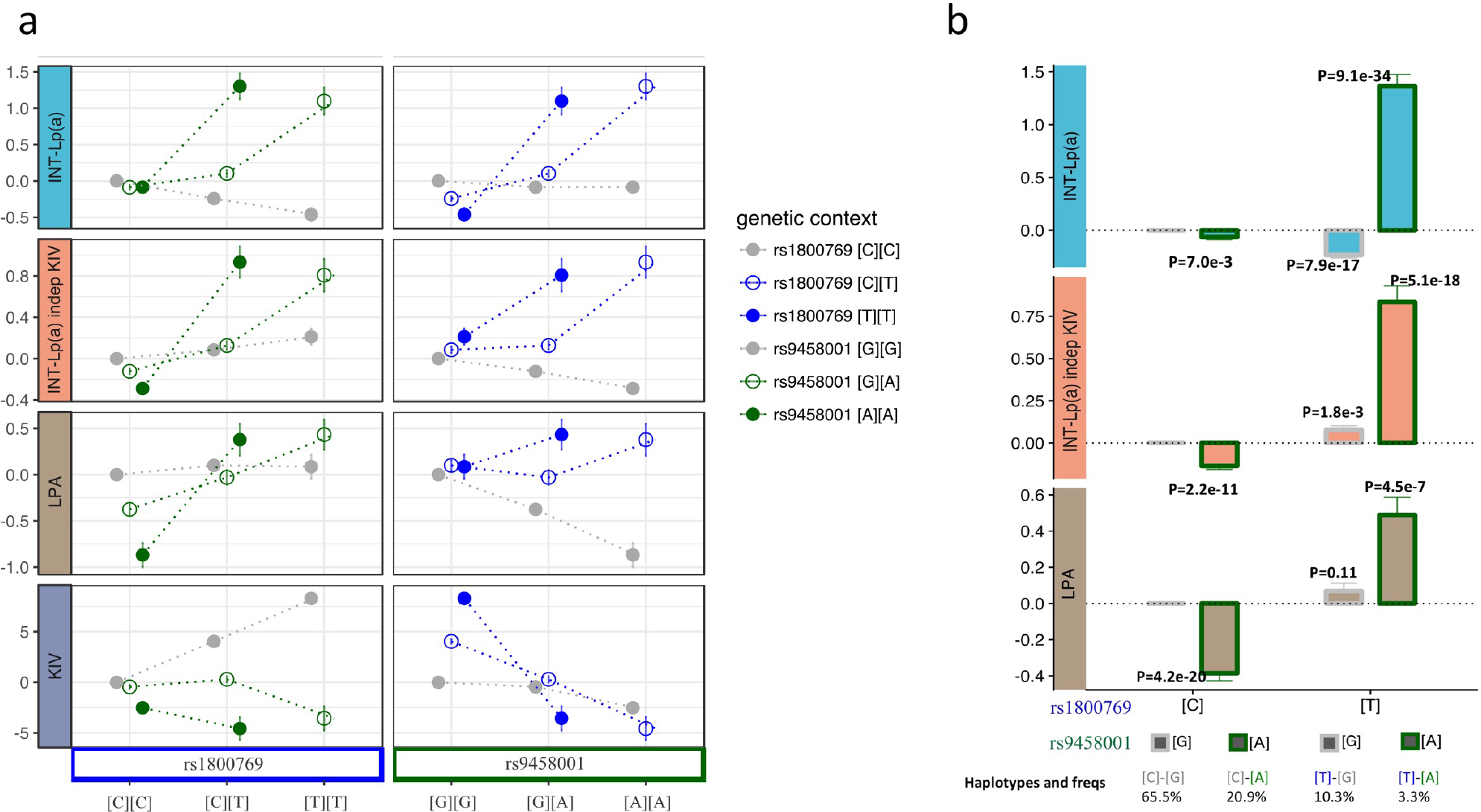
Statistical interaction between SNP effects on circulating Lp(a) level and apo(a) expression activity. **a.** Relative effect sizes (with reference to the genotype [C][C]-[G][G]) are displayed for 8 genotype subgroups ([TT]-[AA] carriers were not found in KORA) respectively for inverse-normal transformed total Lp(a) level (INT-Lp(a), light blue), inverse-normal transformed total Lp(a) level independent of the KIV size (INT-Lp(a) indep KIV, salmon), *LPA* mRNA expression in liver (LPA, brown), and KIV size of the dominantly-expressed apo(a) isoform (KIV, purple). **b.** Relative effect sizes for monoallelic-wise 4 haplotypes are displayed in bar graph with the error bars representing the standard errors, respectively on inverse normal transformed total Lp(a) level (light blue), the total Lp(a) level but with adjustment of the prevalent KIV size in the same sample (salmon), and the LPA mRNA expression in liver (brown). The statistics for the original Lp(a) level without transformation are provided in Supplementary Table 12 and 13.

Circulating Lp(a) levels show high variation in the European population^31^ and are modulated by at least two independent mechanisms. Firstly, they are inversely correlated with the number of Kringle IV type 2 repeats (KIV-2 copy number variation [CNV])^27,28^, which accounts for about 18% of the variability in Western Europeans^32^. However, individuals with the same number of KIV-2 CNV may still differ up to 200- fold with respect to Lp(a) levels^27,28^, suggesting also transcriptional mechanisms. Indeed, in the KORA population we observed an association of the same interaction with KIV-2 CNV (P=1.2×10^−30^) (Supplementary Table 6, Supplementary Note IV). Notably, the risky allele combination [T-A] was accompanied by predominance of shorter KIV-2 CNV variants (i.e., 20-22 repeats) (Fig 3, Supplementary Fig 9), which facilitate Lp(a) release from liver cells. However, both SNPs are in minimal LD with the reported 61 KIV-2 CNV representable variants or the 3 independent modifier variants that influenced the relationship between KIV-2 CNV and Lp(a) cholesterol^32^ (Supplementary Table 9). More importantly, interaction of the two SNPs regarding Lp(a) levels remained highly significant after adjustment for the KIV-2 CNV (P=2.6×10^−11^) (Fig 3, Supplementary Table 6). Therefore, we extended our investigation to apo(a) mRNA expression in liver tissue (Methods, STARNET study, n=522), where *LPA* is transcribed and further assembled to Lp(a). Interestingly, a significant interaction between the two SNPs was found again (P=1.4×10^−8^) and the effects on apo(a) mRNA expression and circulating Lp(a) levels correlated for various genotype subgroups (Fig 3, Supplementary Table 6, Supplementary Fig 8). This suggests that differential gene expression activity underlies a large component of the statistical interaction related to the two SNPs.

Finally, we wish to propose a hypothetical molecular mechanism of epistasis in that the expression activity of the *LPA* gene is determined by the two SNPs co-regulating the enhancer-promoter interaction (Supplementary Fig 10). On the one hand, *in vitro* studies have shown that the rs1800769[T] allele (which lies within at the apo(a) promoter region, Supplementary Table 10) leads to a higher apo(a) transcriptional activity^33^. On the other hand, the rs9458001[A] allele (which lies upstream of the apo(a) promoter) affects multiple binding motifs (Supplementary Table 10), suggesting alteration in enhancer co-activators. Moreover, given that CTCF is a critical regulator of context-dependent enhancer-promoter interaction^34–36^, we investigated the three CTCF-DNA binding sites spanning from rs1800769 and rs9458001 (Supplementary Fig 10, Supplementary Note V). Interestingly, rs1800769[T] corresponds to higher local CTCF-DNA binding affinity at position 1 (Supplementary Table 10), whereas rs9458001[A] facilitates CTCF-DNA binding at position 2 (via a decrease in local CpG methylation^37^, Methods, Supplementary Table 11), which in turn changes the CTCF-cohesin topological structure (Supplementary Fig 10). Combinatorially, the two SNPs may form context-dependent promoter-enhancer interactions in four scenarios that finally lead to differential gene expressions on a large magnitude (Supplementary Fig 10). A note of caution needs to be added, however, since the discussed SNP-SNP interaction is just one of many possible interactions as there are other SNPs at the locus that are in LD with the ones that we discovered.

An inherent challenge in testing for epistasis of nearby SNPs, even if they are in very low LD, is to discriminate interacting SNPs from SNPs representing a specific haplotype. In our case, it seems that the profound biological effects of the interacting pair were amplified in that the genotype combination leading to the highest transcriptional activity is found on a haplotype that contains a low number of KIV-2 CNV repeats, which by independent mechanisms may lead to higher Lp(a) serum levels. However, Lp(a) serum concentrations adjusted for the KIV-2 CNV and apo(a) expression levels were both strongly affected by the interacting SNP pair, which strongly argues – when taken together with other genetic and molecular data (Supplementary Note VI) – for a true epistatic effect.

In summary, we have identified for the first time a SNP-pair at the LPA locus that epistatically affects CAD susceptibility via large-scale statistical interaction analyses. Consistent effects were validated in multiple cohorts, across different cardiovascular traits and intermediary risk factors and traced all the way to apo(a) expression in the liver. The data reemphasize profound biological effects of massively elevated Lp(a) levels in increasing cardiovascular risks. We propose a hypothetical mechanism underlying epistasis of the two SNPs in coordinating gene regulation, and call for further research of epistasis in complex disease.

## Online Methods

### CAD Case-control studies

Individual level genotypes were obtained from ten CAD case-control studies. Most individuals were collected from Germany and have been published before: the German Myocardial Infarction Family Studies (GerMIFS) I^17^, II^18^, III (KORA)^19^, IV^16^, V^20^, VI^2^, VII and the LUdwigshafen RIsk and Cardiovascular Health Study (LURIC)^38^; from Germany and France: Cardiogenics; from England: Wellcome Trust Case Control Consortium (WTCCC)^21,22^; and from France, Italy and the United States: Myocardial Infarction Genetics Consortium (MIGen)^22,23^. GerMIFS VII represents CAD cases with early and severe onset of disease from the Deutsches Herzzentrum München and population-based controls from the Heinz-Nixdorf-Recall (HNR) Study^39^.

Data for WTCCC were obtained via the Leducq network “CADgenomics” (https://www.fondationleducq.org/network/understanding-coronary-artery-disease-genes/). Data for MIGen were obtained via the database of Genotypes And Phenotypes (dbgap)^40^ (project ID #49717-3). All subjects in all studies were of European origin and gave written informed consent before participating. All individuals provided informed consent that specifically addresses that the materials will be used for studying the effect of genetic variants on coronary risk. All respective studies have obtained IRB approval from their local Ethical Committees. Ascertainment and assessment methods CAD of each study is provided in the corresponding publications. Sample size in Supplementary Table 1, and the genotype processing procedures including QC and imputation are provided in Supplementary Notes II.

### UK Biobank

The UK Biobank project (http://www.ukbiobank.ac.uk) is a large prospective cohort study of ∼500,000 individuals from across the United Kingdom, aged 40-69 years at recruitment^41^. Following informed consent, a rich variety of phenotypic and health-related information was collected for each participant, making the resource unprecedented in its size and scope. In addition to self-reported information, including basic demographic data, dietary and exercise habits, multiple physical, cognitive and biochemical measurements were obtained. However, biochemical data for each of the four main blood lipids were not available at the time of analyses. UK Biobank participants are being followed up through electronically linked health-related records. Health-related outcome records include death notifications and cancer diagnoses through linkage to national death and cancer registries, and hospital inpatient episode statistics, which contain coded data on admissions, operations and procedures (primary and secondary). In this study, CAD cases were defined using the “HARD” criteria^2^ as individuals with fatal or nonfatal myocardial infarction (MI), percutaneous transluminal coronary angioplasty (PTCA) or coronary artery bypass grafting (CABG). Peripheral arterial disease (PAD) cases were defined as self-reported history of PAD or leg claudication/ intermittent claudication, or hospitalization or death due to ICD9-443.9, 444, ICD10-I73.9,I74. Aortic valve stenosis cases were defined as self-reported history of aortic stenosis, or hospitalization or death due to ICD9-424.1, ICD10-I35.0.

### KORA F3/F4 studies

Individual-level genotypes were obtained from Augsburg population studies from Germany^30^: KORA F3 and KORA F4^42,43^. The KORA F3 study, conducted in the years 2004/05, is a population-based sample from the general population living in the region of Augsburg, Southern Germany, which has evolved from the WHO MONICA study (Monitoring of Trends and Determinants of Cardiovascular Disease). The KORA F4 survey is an independent non-overlapping sample drawn from the same population in the years 2006/08. The lipid measurements include total Lp(a) levels, and the number of kringle repeats of the Lp(a) protein, determined by Western blotting^24^. Apo(a) isoforms were determined by quantitative analysis of the Western blots. For each analysis, which included isoforms/KIV-2 copy number variation (CNV) repeats, the predominantly expressed isoform for each person was used. More detailed information is given in Supplementary Note IV.

### STARNET Study

RNAseq have been generated from liver in a total of 522 CABG CAD patients from the Stockholm-Tartu Reverse Network Engineering Task (STARNET) study^44^. All patients were Caucasian (30% females), 27% had diabetes, 77% had hypertension, 68% had hyperlipidemia, and 37% had a myocardial infarction before age 60. Patients diagnosed with CAD who were eligible for open-thorax surgery at the Department of Cardiac Surgery, Tartu University Hospital were enrolled. Informed consent was obtained from all subjects (Ethics Approvals Dnr 154/7 and 188/M-12). Blood DNA Genotyping was done using the Illumina Infinium assay^45^, and the results were analyzed with GenomeStudio 2011.1 (Illumina). A 2–3 cm incision in the diaphragm was made to access the peritoneal cavity. This incision was placed to enable direct access to the outer edge of the left lateral liver lobe, from which a 3–5 mm^3^ biopsy was obtained. The liver incision was sutured to control any bleeding. Samples RIN scores of > 7 were accepted and sequenced on Illumina HiSeq with single-end at read length 50 or 100 base pairs. DNA and RNA qualities were assessed with the Agilent 2100 Bioanalyzer system (Agilent Technologies, Palo Alto, CA). Detailed procedures for genotype and RNA processing, including QC and imputation are provided in Franzén et al 2016^44^.

### Broad-sense CAD susceptibility regions

We focused our analysis on loci with previous evidence of genome-wide association with CAD in order to restrict the number of variants for testing of statistical epistasis with the aim to enhance speed and the likelihood of positive finding. We collected the lead SNPs from the 56 published CAD susceptibility loci^16,22^, and decided on ±500kb as a balanced threshold for the flanking range surrounding the known loci (i.e. the broad-sense CAD susceptibility regions), which could meanwhile maximize the heritability covered by the regions and minimize the computational burden. Indeed, variance explained by the lead SNPs only achieved 46% proportion to that could be explained by including their flanking ±500kb regions together (Supplementary Note I). Altogether there are 8,068 SNPs with pairwise r^2^ < 0.5 located in the broad-sense CAD susceptibility regions.

### Statistical interaction analysis for CAD

We used the general framework of detecting statistical epistasis in quantitative genetics as proposed by Hansen and Wagner^46^, and as a pilot study focused specifically on the pairwise epistasis between two loci (SNPs). Formally (Eq. (1)), the case-control status of each CAD individual was taken as a quantitative dependent variable *y* in a linear model, with s*np1* and *snp2* representing the encoded individual genotype for each SNP, respectively. The genotype encoding each variant included four possible models, i.e., dosage (minor allele copy counts; absence as zero), dominant (presence of at least one minor allele copy counts as one; absence as zero), heterozygous (presence of only one minor allele counts as one; otherwise zero), and recessive (two copies of the minor allele counts as one; otherwise zero). Independent variables are *snp1*, *snp2*, and their dot product. Regression was performed for each combination of SNP-pair. The corresponding regression coefficients were estimated with *b*_*1*_, reporting the main effect of the coded variable of *snp1*, *b*_*2*_ reporting the main effect of the coded variable of *snp2*, and *b*_*int*_ for the interaction effect (epistasis coefficient) of the coded variables (*snp1* and *snp2*). In this way, the epistasis coefficient *b*_*int*_ indicates the directionality and quantifies the strength of the effect between both loci (*snp1* and *snp2*).

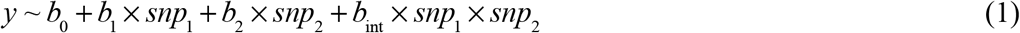

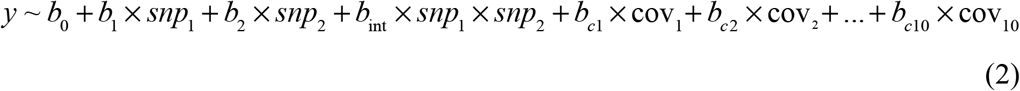

We started exploration of statistical epistasis with all available SNPs located in the broad-sense CAD susceptibility regions across the genome but with LD redundancy pruned to pairwise r^2^ < 0.5 (n_SNP_LDpruned_ =8,068). The statistical interaction calculations were done on this set of SNPs with a low LD structure, with *I* – primary filtering of potential candidates, and *II* - screening and final confirmation (Supplementary Fig 1). Step *I* aimed the fast speed identification of potential significant interaction terms, as well as their respective genotype models, with the assistance of GLIDE GPU computation tool^47^ (n_tests_of_each_model_ = n_SNP_LDpruned_ × (n_SNP_LDpruned_ − 1) / 2 = 32,542,278; n_genotype_models_ = 4 ×4 = 16). General linear regression was performed with the basic model of epistasis (Eq.(1)). A loose and arbitrary significance level was applied (p < 1e-8) for primary filtering with the assumption that if there exists true epistasis between two lead SNPs, loose signals should be detectable between the SNPs within the corresponding LD block. Step *II* included the fine-mapping of the candidate SNP pairs to screen out the pairs with the strongest signal amongst the multiples SNPs in the same LD block. LD-based clumping was performed via PLINK^48^ (v1.90b3.42) to determine the total number of LD independent SNPs (n_SNP_indep_ = 4,654) resulted from step *I*, which was then used to calculate the final significance level with Bonferroni correction 0.05 ⁄ (n_SNP_indep_ × (n_SNP_indep_ − 1) / 2) = 4.6178e^−9^. In total a set of n_SNP_all_ =7,579 variants spanning across the complete block, were further investigated for fine-mapping using logistic regression, performed in R (n_tests_of_given_model_ = n_SNP_all_ × (n_SNP_all_ − 1) / 2 = 28,716,831). Here variants were encoded in the most significant genetic models resulting from Step *I*, and the equation was extended (Eq. (2)) to correct for population stratification and employing a logistic model for the dependent variable. Population structures for each cohort were captured in the genotyped data with multidimensional scaling (MDS) analysis of the identity-by-state (IBS) matrix, computed via PLINK^48^ (v1.90b3.42).

In the discovery phase, the same epistasis testing was performed based on 1000G imputed genotypes in each of the ten CAD case-control studies separately with the adjustment of top 10 MDS components, and then fixed-effect meta-analysis to estimate the overall effect size and standard error. The final epistasis pair of interest was then reanalyzed in the same study with HRC imputation to enable a more complete coverage of the region of interest in all 10 cohorts. This gave consistent results, showing that the finding obtained was not dependent on the imputation panel used. Additionally, individual-level genotypes of two independent cohorts, GerMIFSVII and UK Biobank (Supplementary Table 1), were utilized as independent replications for the top lead SNP pair. Analysis based on GerMIFSVII was adjusted with covariates of top 5 multiple dimensions. Analyses based on UK Biobank data were adjusted with covariates of top 5 principal components, age, gender and genotype platform.

### Prioritizing candidate SNP pairs of epistasis of CAD

After the detection of all SNP pairs showing statistically significant epistatic effects on the risk of CAD, we prioritized the candidate pairs based on the following rules of thumb and selected the top candidate with the consideration of further functional investigations.

Firstly, we picked up SNP pairs that were highly replicated, as previous statistical epistasis explorations have been frequently blamed for low replication^3,49^. Here we retained only SNP-pairs, which displayed statistical epistasis both significantly and consistently in at least 8 studies (out of 10) in the discovery data, based on both v1000G and vHRC imputation.

Secondly, when our statistical interaction SNP pairs were located on the same chromosome (i.e., cis-epistasis), we clarified the regional LD structure to confirm the independence of LD between two target SNPs (*snp1* and *snp2*). We filtered out all pairs with LD r^2^ > 0.2 between *snp1* and *snp2* (inter-pair criteria). Moreover, we assumed that if significant and true statistical epistasis effect exists between *snp1* and *snp2*, then at least weak interaction signals should be detectable between all extended SNPs in high LD with each of the two interacting SNPs (respectively all proxies of *snp1* and all proxies of *snp2*), unless either SNP itself is a LD singleton and has limited or no other variants nearby linked with it. Therefore, to maximize the replication of the true positive discovery, we focused on those pairs with both target SNPs as non-LD-singleton. All singletons or SNPs with low LD (r^2^ between 0.2 - 0.5 with any other variants) were not further considered. All SNPs in LD r^2^ > 0.5 were grouped into the same target SNP group (Supplementary Fig 2a) as *snp1* group or *snp2* group (intra-pair criteria), and the SNP pair with the largest effect size for the interaction coefficient was taken as the lead pair (Supplementary Table 3).

Thirdly, to test the independence of the statistical epistasis effect against the effect from a third SNP in the vicinity of *snp1* or *snp2*, or from any known CAD susceptibility loci reported via traditional GWAS approach, conditional analyses were performed in all ten studies followed by meta-analysis. Dosage model was assigned to the third SNP (*snp3)*, extracting from the latest compiled list of 164 known CAD GWAS SNPs^50^, and from all available SNPs in the ±200kb flanking regions of *snp1* and *snp2*. Logistic regression was utilized to estimate the effect of the *b*_*int*_ (epistasis coefficient) but with *snp3* as an additional covariate (Eq. (3)).

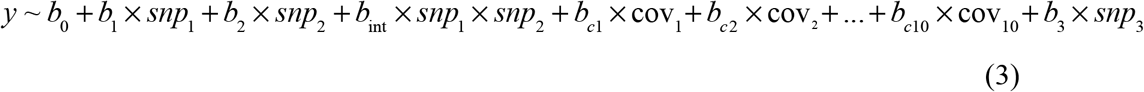

As a result, four pairs displaying consistent effects in at least 8 studies, hit all the defined criteria for statistical epistasis (Supplementary Table 2), with the top pair as two SNPs rs1800769 and rs9458001 on chromosome 6 in dosage-dosage model. The effect sizes were consistent across all studies, except for GerMIFSI due to the unavailability of one of the SNPs (or any proxy) in the pair based on 1000G imputation, but proxies (r^2^ ≥ 0.98) were available in our data based on HRC imputation and showed consistently strong interactions across all ten studies (Supplementary Fig 3). With the aim to statistical-to-biological translation, we annotated the genetic locations of the 8 SNPs in these four candidate pairs (Supplementary Table 2), and found one of the pairs with both SNPs located nearest to a known CAD risk gene – LPA, i.e., the pair of rs1800769 and rs9458001, with the former at the *LPA* promoter and the latter at the upstream intergenic region. We thereafter focused on this top epistasis SNP pair with the highest potential to be biologically relevant for further biological and functional analyses, and argue against *cis*-epistasis being a pure statistical artifact^51^.

### Statistical interaction analysis on intermediate factors

For the lead SNP pair that displayed epistasis effect on the risk of CAD, we further tested for signals on a series of intermediate factors. Regression analyses were performed using R based on the same genetic model (Eq. (1)) but we replaced the dependent variable *y* as the corresponding intermediate factor of our interest. Covariates were added if necessary for each intermediate factor.

Circulating Lp(a) were measured in KORA F3 and F4 studies^42,43^. Due to the highly skewed distribution of Lp(a), inverse normal transformation was applied to the Lp(a) concentration to construct the dependent variable *lpa* (Eq.(4)). Analysis was first performed for KORA F3 and F4 separately. Then meta-analysis was performed to estimate the effects and p-value.

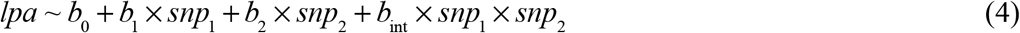

For a sample of 2,831 LURIC individuals, a series of cardiovascular related risk factors were also measured including circulating Lp(a) concentration, in addition to the records of CAD status. Lp(a) was measured in EDTA plasma in the scale of mg/dl. Logistic regression was performed for CAD status, with (Eq.(6)) and without (Eq.(5)) adjustment for the inverse normal transformation Lp(a) to estimate the effect change for the epistasis coefficient *b*_*int*_

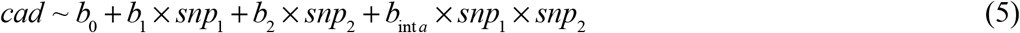

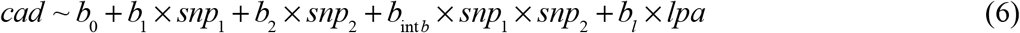

For the epistasis effect on the apo(a) expression activity we applied two approximate measurements: a) the effect on the hepatic LPA-mRNA expression by RNA-seq from STARNET^44^ Study (Eq. (7)); and b) the effect on the total Lp(a) level adjusted by the effect related due to KIV-2 CNV (Eq. (8), Supplementary Note IV). Lp(a) concentration was inverse normal transformed. Analysis was first performed in KORA F3 and F4 separately. Then meta-analysis was performed to estimate the effects and significance.

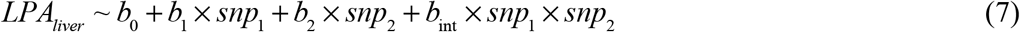

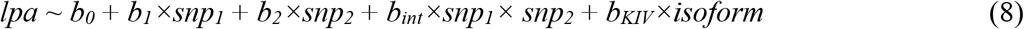

### Relative effect size analysis for genotypes and haplotypes

The effect of interaction term *b*_int_ indicates the deviation of the observed effect compared to the expected combination of effect given the respective effect-allele models of the two SNPs, but does not tell the actual risk difference among subgroups of individuals divided according to the interactive SNP-pair. In order to characterize the manifestation of interaction as to how the genetic effect of one SNP is mutually dependent on the genetic context of the other SNP, we dissected the genetic effect of epistasis by analyzing the effect of two SNPs from two perspectives: in genotypes (e.g., biallelic-wise in 3 * 3 = 9 genotype combinations), and in haplotypes (e.g., monoallelic-wise in 2 * 2 = 4 haplotype) (Supplementary Fig 6). For genotypes, we set the effect for the majority group as the baseline for comparison, and performed multivariate regression with 9 genotype combination categories, to estimate the relative effect change of each group on multiple levels (Eq. (9)), including CAD odds ratio, total Lp(a) level, LPA mRNA expression in liver, and isoform-independent Lp(a) level, based on the corresponding data.

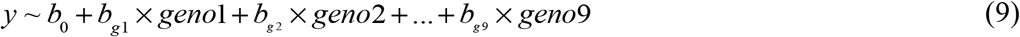

Indeed, monoallelic-wise haplotype would be more direct reflection of the genetic context. Given that only genotypes, rather than haplotypes, are directly measured and observed in genotyping arrays, we inferred estimated haplotypes via linkage phasing of the SNP genotypes for ambiguous (heterozygous at both SNPs) haplotypes using R package *hapassoc*. Firstly, using the genotype data an input a list of possible haplotypes that are compatible with each person’s genotype from the pre-processing function “pre.hapassoc”, which generates dummy samples for uncertain haplotypes, additionally generated with probabilities. Then the haplotype association analyses implemented in the “hapassoc” function were conducted based on generalized linear models using the expectation-maximization (EM) algorithm. Similarly, we set the effect for the majority group as the baseline for comparison, and performed multivariate regression with 4 haplotype categories, to determine the relative effect change of each group on multiple levels (Eq. (10)).

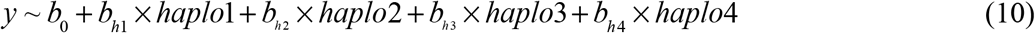

### Methylation QTL analysis

We utilized the resource published by Gaunt et al^37^ containing summary statistics of genetic variants underlying each DNA methylation CpG islands collected at five different time points across the life course from individuals in ALSPAC, with on average about 800 samples for each time point (Supplementary Table 11). Furthermore, we analyzed methylation array data from a subsample of 588 LURIC participants (Illumina EPIC array) measured in blood cells. Raw data were processed following the recommendations from the CKDGen consortium. There were 487 overlapping samples with both genotype and methylation data available that underwent filtering for quality control. All 565 methylation probes in the LPA flanking region (chr6:160363532-161643608, hg19) were extracted. Genotype-methylation associations were performed for rs1800769 and rs9458001, and all non-SNP-overlapping methylation probes (SNP-probe distance >1bp) with p < 0.01 to either SNP were checked. Only rs9458001 was significantly associated with CpG methylation levels, and was reported (Supplementary Table 11). The rs9458001[A] allele was associated with reduced methylation of a CpG island in the region between the two SNPs (∼15kb upstream of rs1800769 and downstream of rs9458001), which was consistent based on the above two independent methylation array analyses.

### Functional annotation

Annovar^52^ was utilized to annotate the genomic location, location relative to the nearby gene, and putative regulatory functions. Haploreg4.1^53^ was utilized to annotate the genomic position, LD structure in the European population, and ENCODE^52^ annotation were integrated to capture possible regulatory elements of promoter, enhancer, and transcriptional factor binding motifs (Supplementary Table 10).

## Acknowledgments

This work was supported by grants from the Fondation Leducq [CADgenomics, 12CVD02], the German Federal Ministry of Education and Research (BMBF) within the framework of ERA-NET on Cardiovascular Disease, Joint Transnational Call 2017 [ERA-CVD: grant JTC2017_21-040], within the framework of target validation [BlockCAD: 16GW0198K], within the framework of the e:Med research and funding concept [AbCD-Net: grant 01ZX1706C and e:AtheroSysMed: grant 01ZX1313A-2014], and the European Union Seventh Framework Programme FP7/2007–2013, under grant agreement no. HEALTH-F2-2013-601456 (CVgenes-at-target). Further grants were received from the Deutsche Forschungsgemeinschaft (DFG) as part of the Sonderforschungsbereich CRC 1123 (B2), from Austrian Science Fund (FWF): Project Number P 266600-B13 to C.L. and the “Genomics of Lipid-associated Disorders (GOLD)” of the “Austrian Genome Research Programme GEN-AU” to F.K. Methylation measurements in LURIC were funded by the 7th Framework Program of the EU [RiskyCAD, grant 305739). C.P.N and N.J.S. are funded by the British Heart Foundation and N.J.S. is a Emeritus NIHR Senior Investigator.

