## Supplementary Text and Figures for "*Cis*-epistasis at the *LPA* locus and risk of coronary artery disease"

### Supplementary Figures

1. **Scheme of statistical interaction test for CAD.**
2. **cis SNP-pair prioritization of statistical interactions of CAD.**
3. **Forest plot showing the epistasis effect on CAD odds ratio for rs1800769 and rs9458001 in each of the ten studies as well as in meta-analysis.**
4. **Forest plot showing the epistasis effect on CAD odds ratio for the other 3 candidate SNP pairs in each of the meta-analysis in the discovery stage.**
5. **Bubble plot showing the signal irrelevance between GWAS and statistical interaction for the lead pair rs1800769-rs9458001.**
6. **Graphical illustration the concept of epistasis through genetic context in 9 genotypes and 4 haplotypes.**
7. **rs1800769-rs9458001 combined effect in comparison with the single dosage effects.**
8. **Demonstration of epistasis between rs1800769 and rs9458001 on CAD and its intermediate factors.**
9. **Distribution of KIV CNV numbers for haplotype rs1800769-rs9458001 [T-A] carriers and non-carriers.**
10. **Hypothetical co-regulation of LPA gene expression by rs1800769 and rs9458001.**
11. **Increased variance explained with physically expansion around the known CAD lead SNPs reported from GWAS studies.**
12. **Relative effect sizes for subgroups of individuals with different genotype context given the other 3 candidate SNP pairs.**
13. **Overview of functional annotations for the genomic region in the vicinity of rs1800769 and rs9458001.**

### Supplementary Notes

**I. Increased variance explained with physically expansion around the known CAD lead SNPs reported from GWAS studies.**

**II. Genotype processing for 10 CAD case-control studies**

**III. Clinical significance of epistasis**

**IV. Size of Apo(a) isoforms and KIV-2 CNV numbers**

**V. Hypothesized mechanism of epistasis**

**VI. Evidence against haplotype effect**

### Supplementary Figures

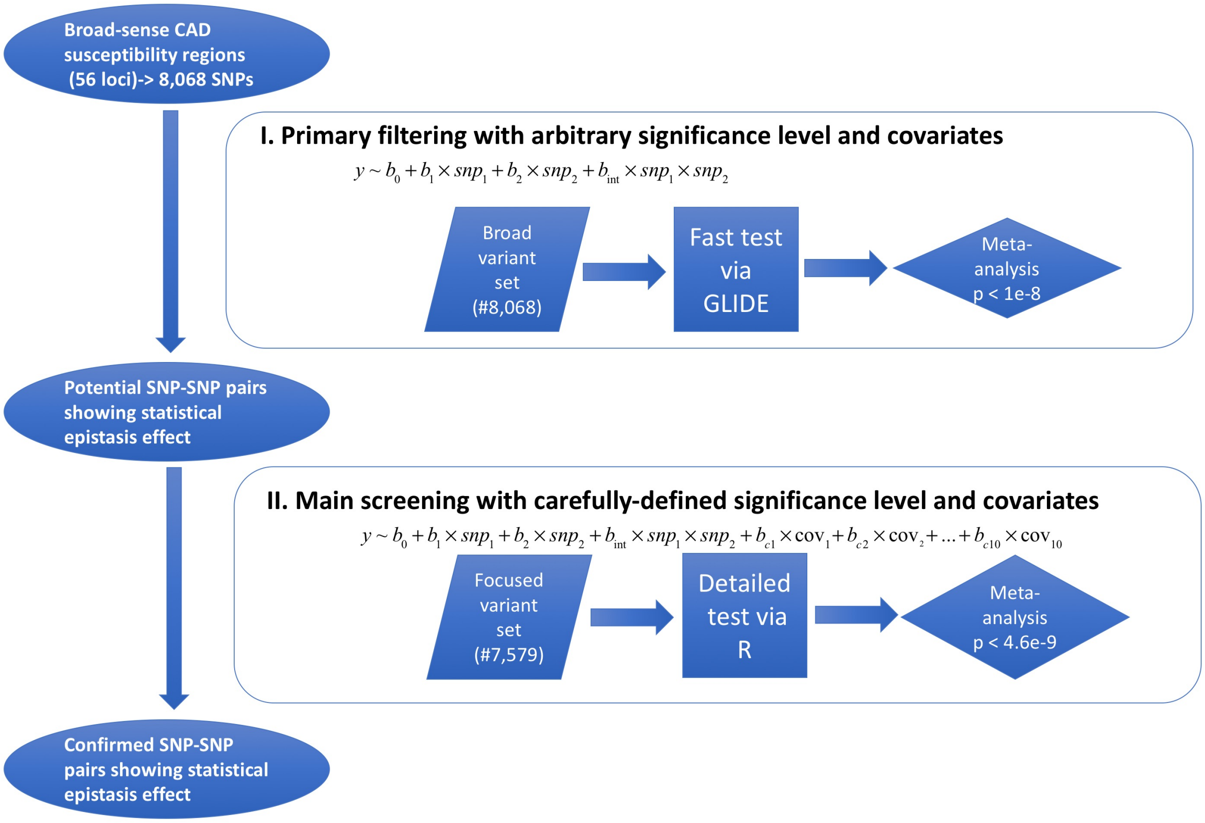


**Supplementary Figure 1**

**Scheme of statistical interaction test for CAD.**

We started exploration of statistical epistasis with all available SNPs located in the broad-sense CAD susceptibility regions across the genome but with LD redundancy pruned to pairwise $r^2 < 0.5$ ($n_{SNP\_LDpruned}$ =8,068). Statistical analysis was conducted with two steps: *I* – primary filtering of potential candidates, and *II* - screening and final confirmation. Step *I* aimed the fast speed identification of potential significant interaction terms, as well as their respective genotype models, with the assistance of GLIDE GPU computation tool[1] ($n_{tests\_of\_each\_model} = n_{SNP\_LDpruned} \times (n_{SNP\_LDpruned} - 1) / 2 = 32{,}542{,}278$; $n_{genotype\_models} = 4 \times 4 = 16$). General linear regression was performed with the basic model of epistasis. A loose and arbitrary significance level was applied ($p < 1e-8$) for primary filtering with the assumption that if true epistasis exists between two lead SNPs then loose signals should be detectable between the SNPs within the corresponding LD block. Step *II* included the fine-mapping of the candidate SNP pairs to screen out the pairs with the strongest signal amongst the multiples SNPs in the same LD block. LD-based clumping was performed using PLINK[2] (v1.90b3.42) to determine the total number of independent SNPs ($n_{SNP\_indep}$ = 4,654) resulting from step *I*, which was then used to calculate the final significance level with Bonferroni correction $0.05 / (n_{SNP\_indep} \times (n_{SNP\_indep} - 1) / 2) = 4.6178e^{-9}$. In total, a set of $n_{SNP\_all}$ =7,579 variants spanning across the complete block were further investigated for fine-mapping using logistic regression, performed in R ($n_{tests\_of\_given\_model} = n_{SNP\_all} \times (n_{SNP\_all} - 1) / 2$ = 28,716,831). Here variants were encoded in the most significant genetic models resulted from Step *I*, and the equation was extended to correct for population stratification. Population structures for each cohort were captured in the genotyped data with multidimensional scaling (MDS) analysis of the identity-by-state (IBS) matrix, computed using PLINK[2] (v1.90b3.42). Statistical epistasis testing was performed in each study separately, and then fixed-effect meta-analysis to estimate the overall effect size, standard error, and p-value. No significant heterogeneity could be observed in the meta-analyses, for this fixed–effect results are presented here.

a

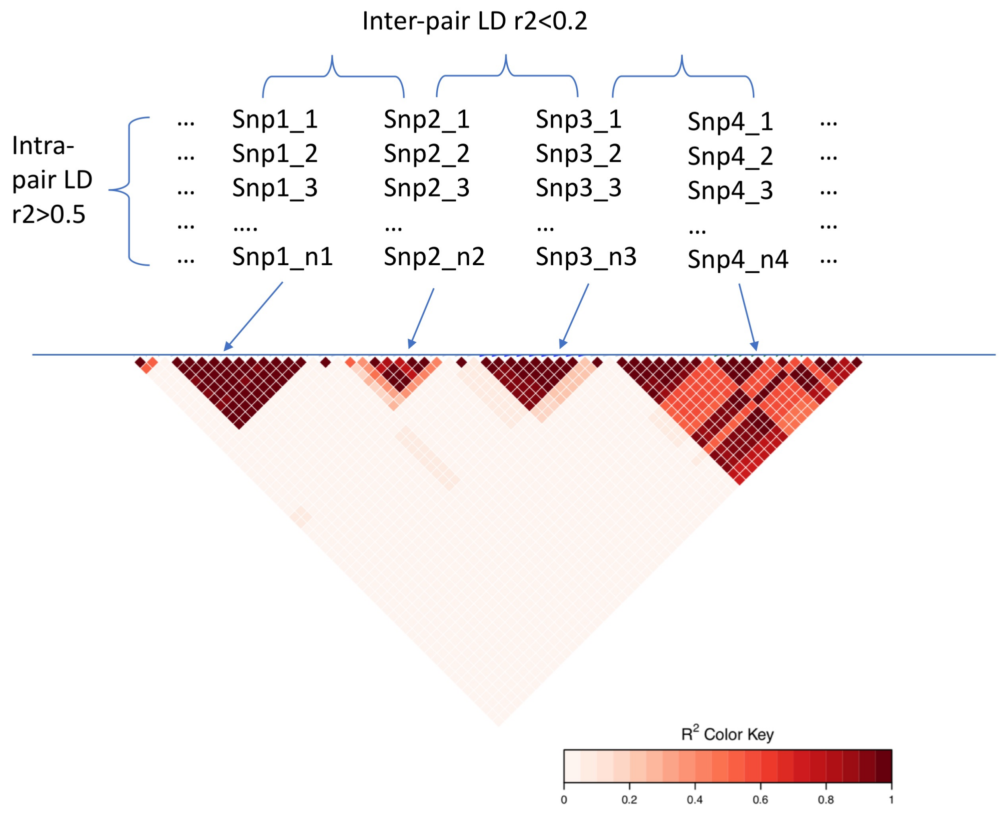

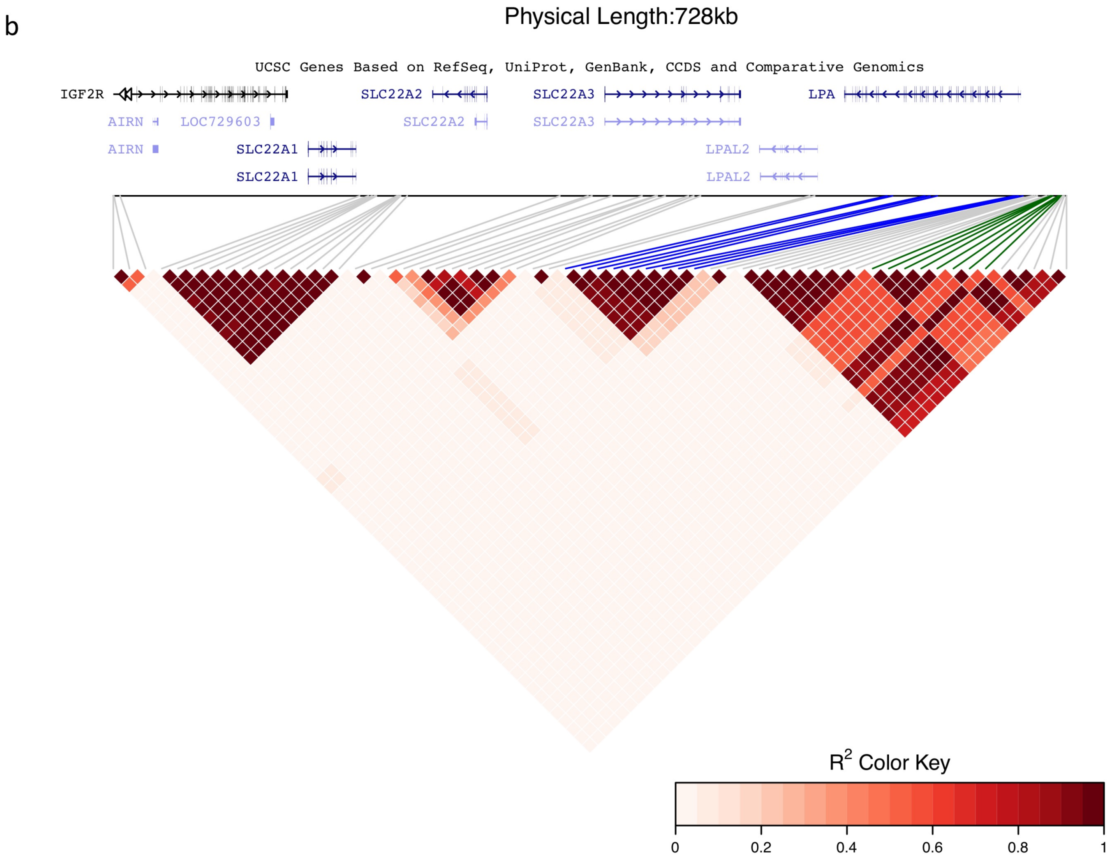


**Supplementary Figure 2**

**cis SNP-pair prioritization of statistical interactions of CAD.**

LD heatmap showing distinct LD blocks along the genome in the vicinity region. Panel **a.** characterizes the methods of defining statistical cis-epistasis. Statistical epistasis pairs were chosen between two non-LD singleton lead SNPs. LD-independent blocks were identified for all variants in the vicinity region. Those with a LD of $r^2 > 0.5$ were grouped into the same SNP bunch, while those with $r^2 < 0.2$ were considered as distinct independent LD blocks. LD singleton (with weak association $r^2$ between 0.2 - 0.5 with any other) were not of our interest for the consideration of replication and interpretation reason. Panel **b**. highlights the genome position of our lead pair of interest. It was selected as the one at the LPA locus, especially interesting, as this gene is a known risk factor of CAD. rs1800769 and rs9458001 are in distinct LD groups. rs1800769 and several of its highly correlated proxy SNPs are marked in blue; rs9458001 and its corresponding proxies are marked in green.

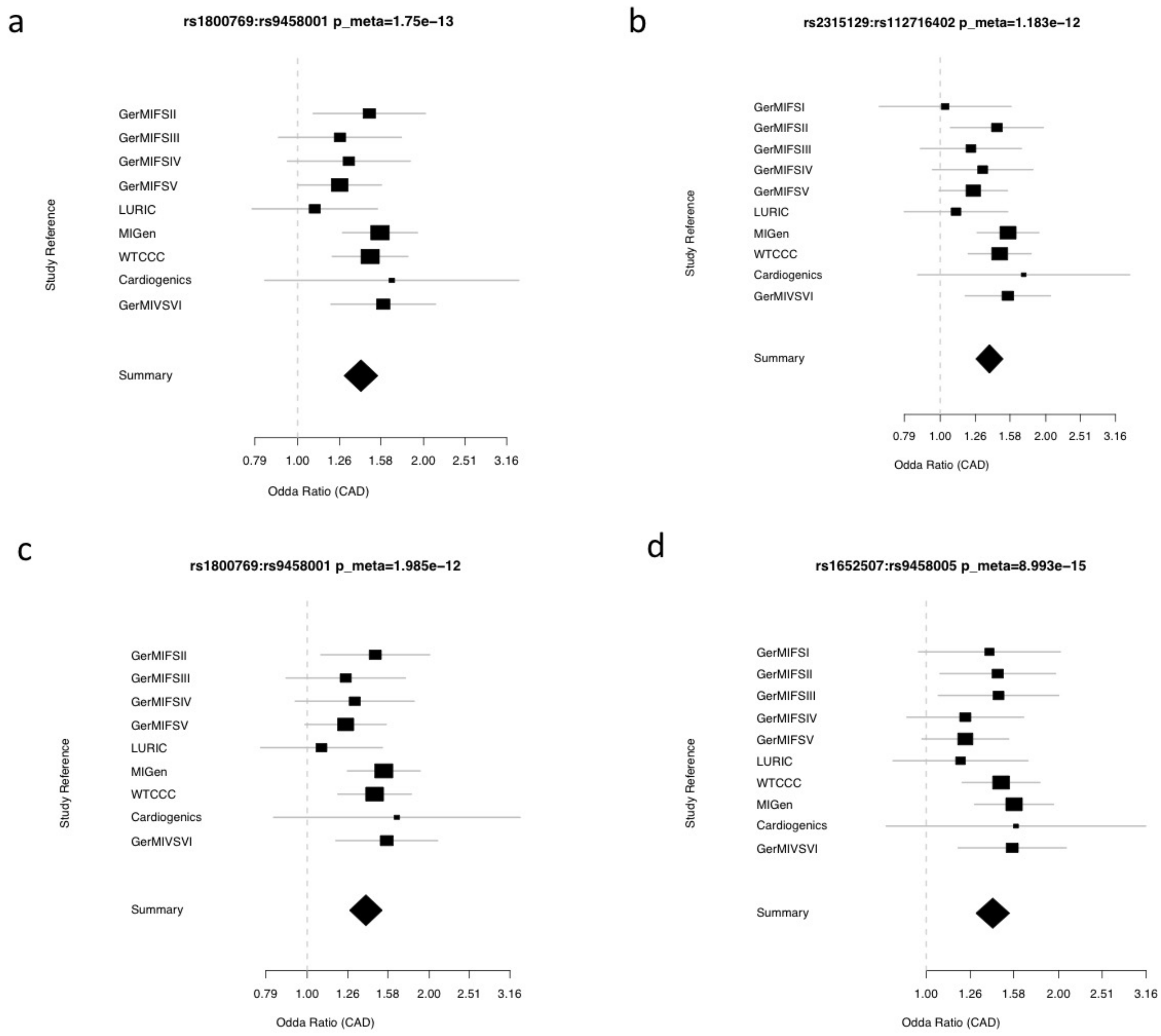


**Supplementary Figure 3**

**Forest plot showing the epistasis effect on CAD odds ratio for rs1800769 and rs9458001 in each of the ten studies as well as in meta-analysis.**

**a.** Lead SNP pair rs1800769 and rs9458001, in the discovery data based on 1000G imputation, as the one with the lowest p-value. (not available in GerMIFSI). **b.** Proxies for the SNP pair rs2315129 (LD $r^2$=0.99 with rs1800769) and rs112716402 (LD $r^2$=0.88 with rs9458001), in the discovery data based on 1000G imputation, as the one available in all studies including GerMIFSI (meta p-value significant, GerMIFSI itself not, but has the same positive trend). **c.** Lead SNP pair rs1800769 and rs9458001, in the discovery data based on 1000G imputation, with the effect conditioned on univariate effect of rs10455872 (which is a top CAD GWAS SNP in the vicinity). **d.** Proxies for the SNP pair rs1652507 (LD $r^2$=0.98 with rs1800769) and rs9458005 (LD $r^2$=1 with rs9458001), in the replication data based on HRC imputation, as the one available in all studies.

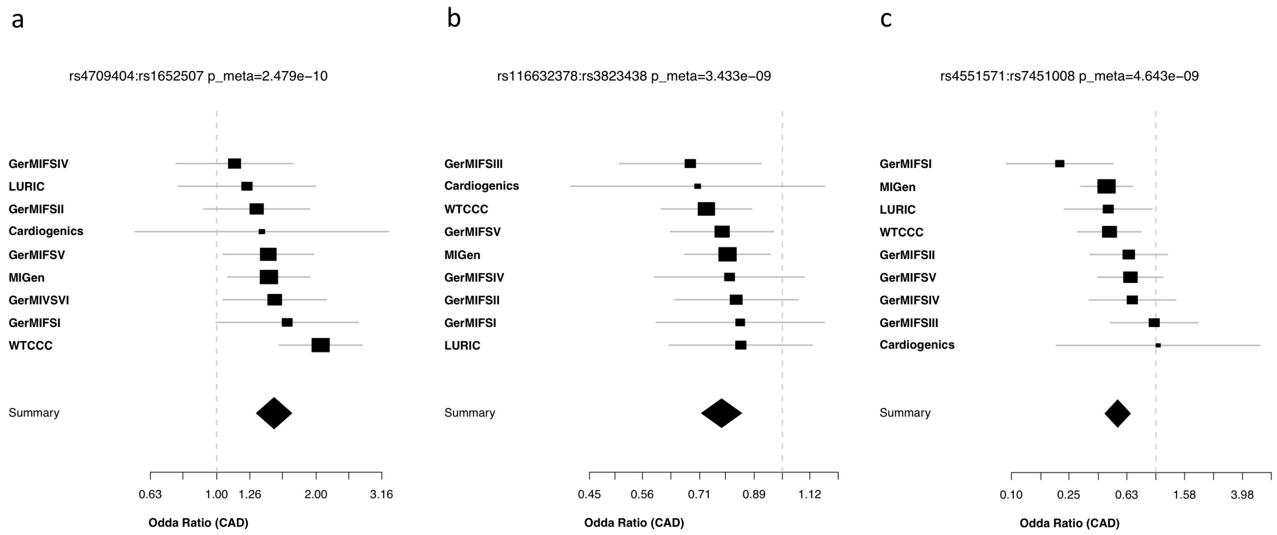


**Supplementary Figure 4**

**Forest plot showing the epistasis effect on CAD odds ratio for rs1800769 and rs9458001 in each of the ten studies, as well as in meta-analysis.**

Square and horizontal line display the estimated odds ratios and 95% confidence interval for each cohort**.** The size of the square is inversely proportional to the standard error of the estimated effect. Below the individual cohorts, a summary diamond depicts the fixed-effects when analyzing all cohorts jointly. **a.** cis SNP pair rs4709404 and rs1652507 in [T] dosage and [C] dosage model. **b.** cis SNP pair rs116632378 and rs3823438 in [T] dosage and [G] dominant model. **c.** trans SNP pair rs4551571 and rs7451008 in [C] dominant and [C] recessive model.

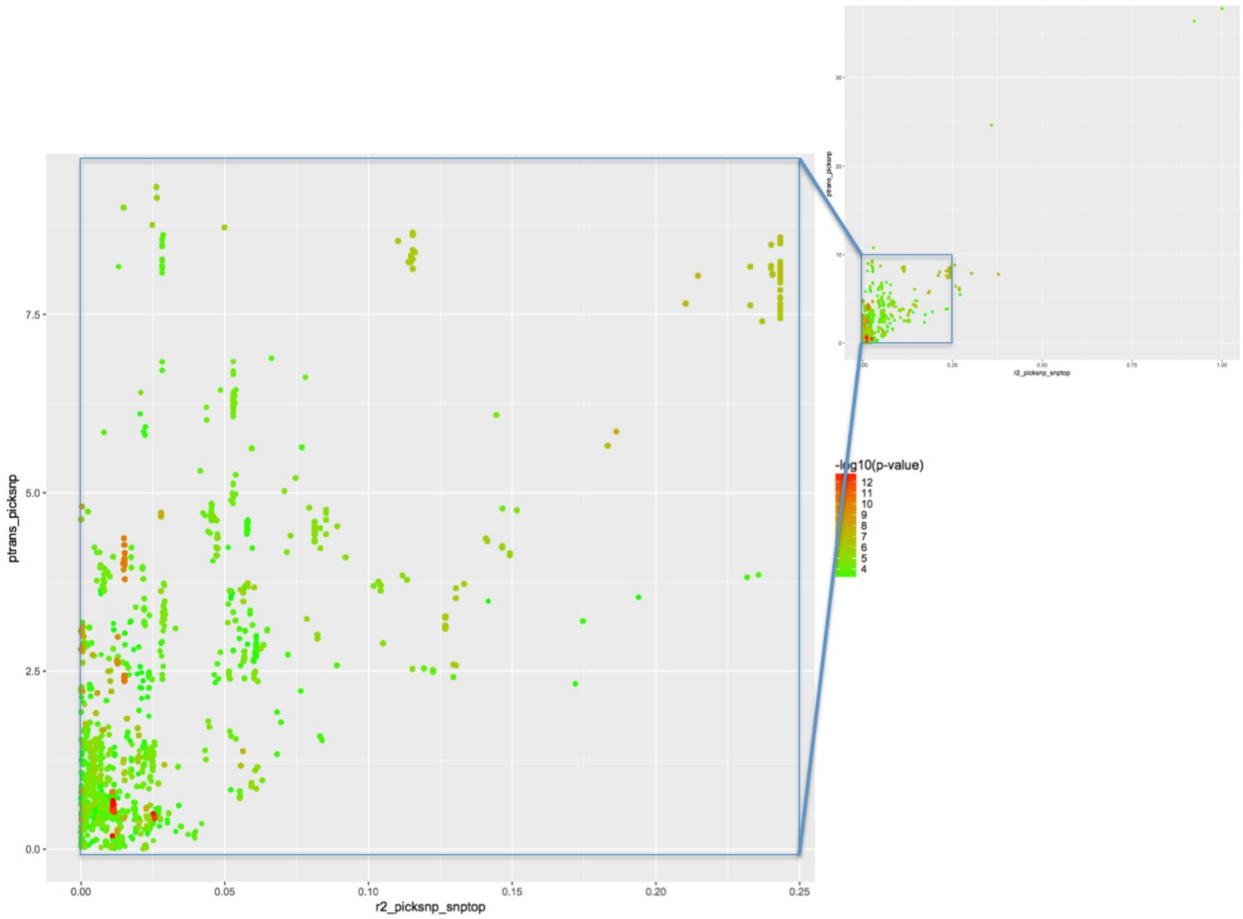


**Supplementary Figure 5**

**Bubble plot showing the signal irrelevance between GWAS and statistical interaction for the lead pair rs1800769-rs9458001.**

For all pairs of SNPs (each pair as a point in the figure) where the statistical interaction $p_{int} < 1e^{-3}$ (meta-analysis based on 10 studies) and with both SNPs in the same LD block as rs1800769 and rs9458001, respectively, we display here the p-values for both signals, i.e., GWAS vs statistical interaction. The y-axis represents the GWAS signal in the measurement of –log10(p-value) for the SNP in one pair whichever with the stronger GWAS signal ("picksnp"). Colors of points, starting with green indicating high p-values ($p_{int} < 1e\text{-}3$) to red indicating low p-values ($p_{int} < 1e\text{-}12$), represent the signal of the statistical interaction in the measurement of –log10(p-value). Moreover, we also display here the linkage disequilibrium (LD) between the "picksnp" and the regional GWAS top SNP - rs10455872) in the measurement of $r^2$ on the x-axis. We can observe that SNP pairs by themselves hav higher GWAS signals (y-axis) do not contribute to more significance (points in reddish color) of the epistasis signal. We can also observe that SNPs by themselves in higher LD with the top GWAS SNP (x-axis) do not contribute to more significance (points in reddish color) of the epistasis signal neither. In short, neither replacing one of the SNP with higher GWAS signal in the same LD block, nor with higher LD with the top GWAS SNP, could increase the significance of the interaction signal. Summary statistics of CAD from Nikpay et al 2015 were utilized to annotate the GWAS p values of all SNPs.

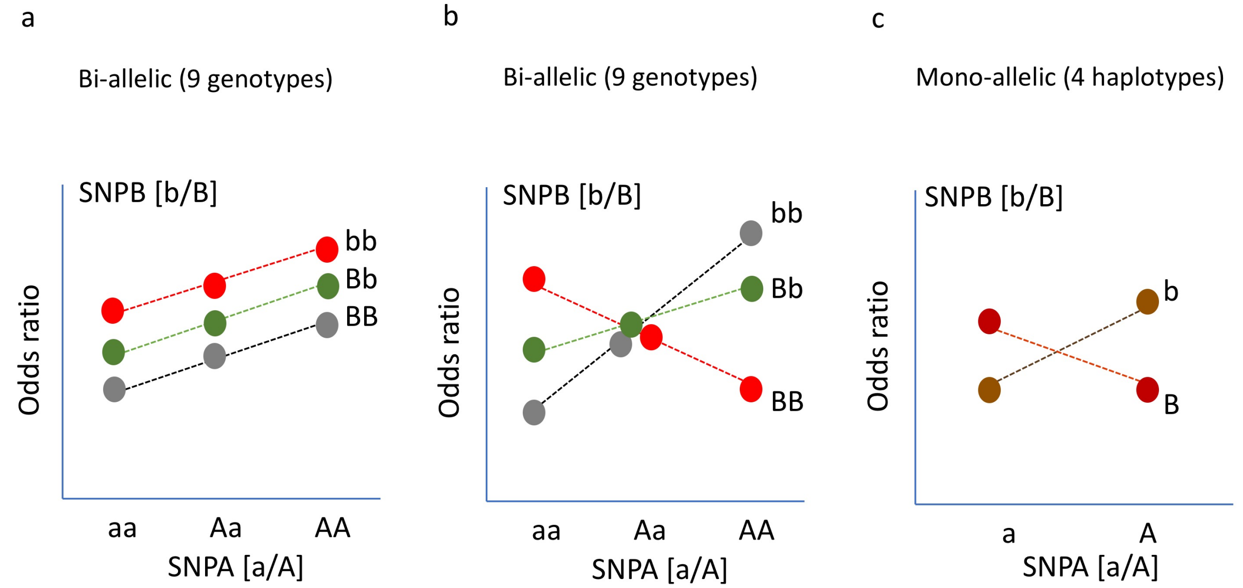


**Supplementary Figure 6**

**Graphical illustration of the concept of epistasis through genetic context in 9 genotypes and 4 haplotypes.**

Toy two-SNP models illustrating the epistatic effect between two SNPs in a dosage-dosage model. Each point represents the effect on a trait of interest (in our case odds ratio of CAD, y-axis) in a biallelic-wise nine genotypes (panel **a** and **b**), and monoallelic-wise four haplotypes (panel **c**) model, respectively. The x-axis represents the three genotypes at one locus (left: homozygotes [aa] for major allele, in the middle: heterozygote [aA], right: homozygotes of minor allele [AA]) while the three dotted lines indicate the genotypes at the interacting locus SNP-B (grey line for homozygotes for major allele [bb], green line for heterozygote [bB], and red line for homozygote of minor allele [BB]). The y-axis displays the mean phenotype scores, respectively.

If no epistasis exists, then the effect size of each SNP is expected to depend linearly or additively only on the number of alleles by that same SNP (regardless of the different genetic context of the other SNP). In other words, the independent SNP effects on the phenotype would just sum up and act in parallel (panel **a**). Whereas in an epistatic scenario the effect of a SNP on a phenotype would dependent on the presence of another locus, which would be indicated by a deviation from additivity, as shown in such a cross shape SNP (graphically crossing-over between the assistance lines) (panel **b**). In practice, as the genotyping process only measures genotypes while each genotype is biallelically from two chromosomes, we performed haplotype phasing to deduce from which chromosome each genotype call came from based on EM estimation (panel **c**).

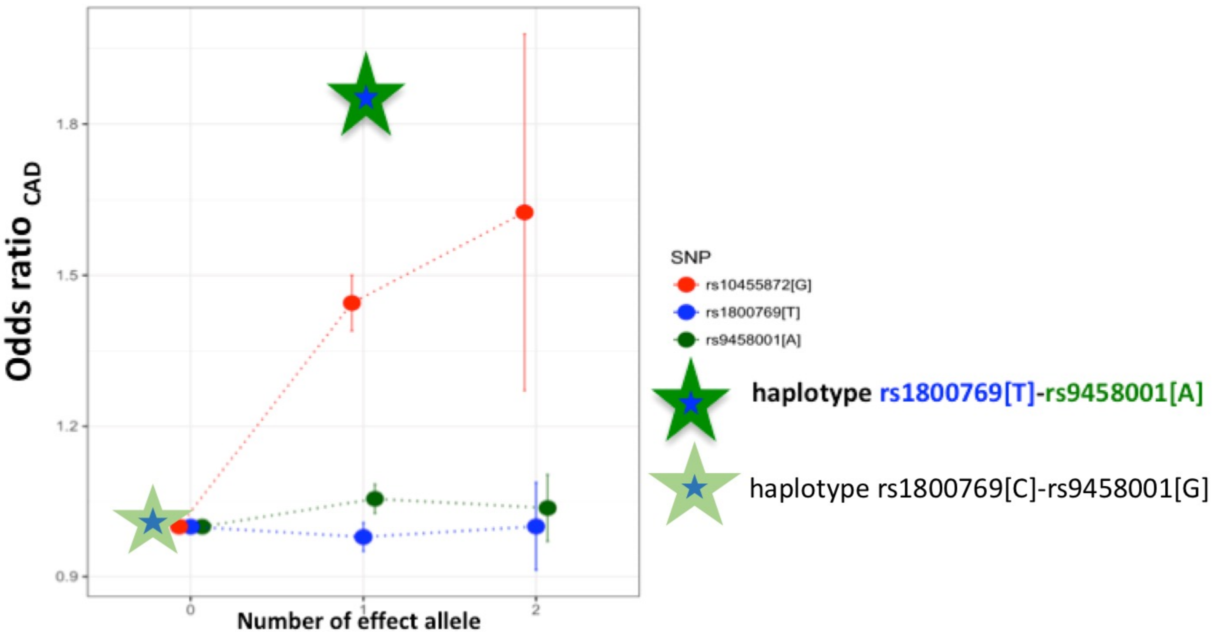


**Supplementary Figure 7**

**rs1800769-rs9458001 combined effect in comparison with the single dosage effects.**

The single and interactive effects of our lead SNPs in the epistasis pair (rs1800769 and rs9458001), as well as the most significant GWAS SNP in the LPA vicinity (rs10455872). Each SNP (rs1800769[T] in blue, rs9458001[A] in green, and rs10455872[G] in red) with its increasing number of effect alleles are aligned along the x-axis. Haplotypes [C-G] and [T-A] (in stars) composed of rs1800769 and rs9458001 are also aligned along the x-axis according to the corresponding number of effect alleles for each SNP are shown as stars in light (reference alleles:

C-G) and dark green (effective alleles: T-A). The y-axis displays the relative odds ratio of CAD for each genotype or haplotype compared to the reference group (for each SNP, the genotype with no effect allele is set as reference; similarly, haplotype [C-G], with no effect allele from both SNPs, is set as reference). It is observed that despite the individually trivial additive effect sizes for rs1800769 and rs9458001, the haplotype that composes both effect alleles from the two SNP (i.e., [T-A]) displays an extremely high risk of CAD (odds ratio =1.84), even larger than the so-far reported genome-wide significant CAD lead SNP rs10455872.

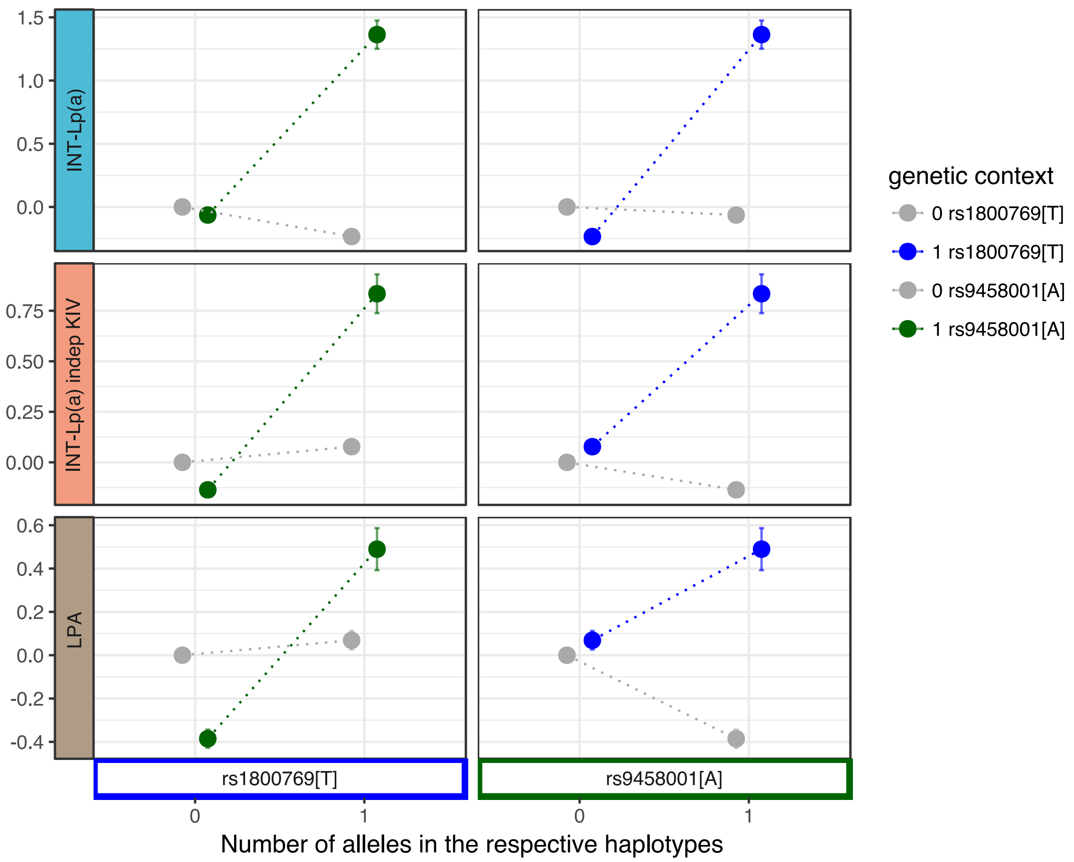


**Supplementary Figure 8**

**Demonstration of epistasis between rs1800769 and rs9458001 on CAD and its intermediate factors.**

Relative effect sizes and standard errors are displayed in monoallelic-wise 4 haplotypes, with the reference haplotype set as rs1800769[C]-rs9458001[G]. The left-hand side shows haplotypes with rs1800769[T] from absence '0' to presence'1' ([C]->[T], labelled in blue along x-axis) against the genetic context of (in co-presence with) of rs9458001[G] (along with the grey dotted line and points as viewing assistance), or against the genetic context of (in co-presence with) of rs9458001[A] (along with the green dotted line and points as viewing assistance). The right-hand side shows vice versa, with rs9458001[A] from absence ‘0’ to presence ‘1’ ([G]->[A], labelled in green along x-axis), against the genetic context of (in co-presence with) of rs1800769 [C], and [T], respectively. The y-axes from panels up to down correspond to the measurement of effect on the inverse normal transformed total circulating Lp(a) level (INT-Lp(a), light blue), the inverse normal transformed total Lp(a) level adjusted with the effect from the KIV-CNV size (INT-Lp(a) indep KIV, salmon), and the LPA gene expression activity in the measurement of LPA gene expression in liver (LPA, brown).

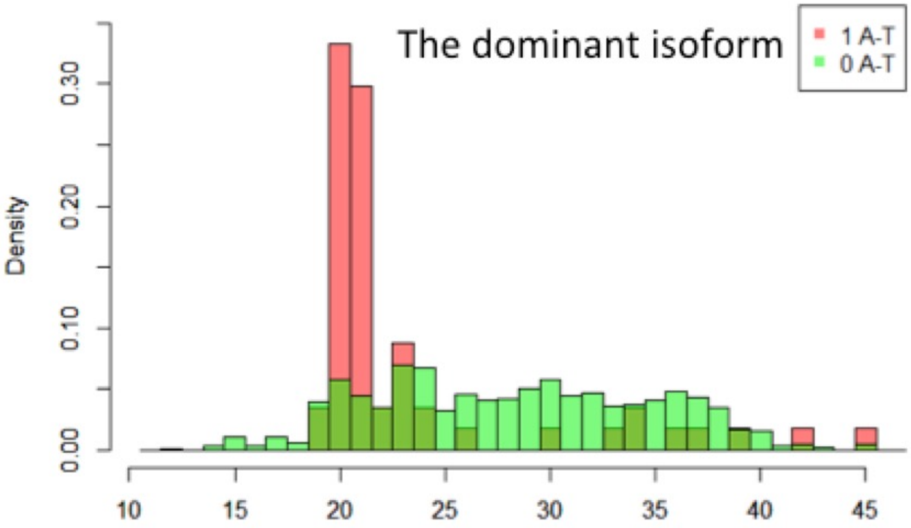


**Supplementary Figure 9**

**Distribution of KIV CNV numbers for haplotype rs1800769-rs9458001 [T-A] carriers and non-carriers.**

For 5,953 KORA individuals, KIV CNV repeat numbers were scrutinized from the Western blot for both the predominantly expressed and weakly expressed apo(a) isoform. Individuals were divided into haplotype [T-A] carriers and non-carriers based on their genotypes for those whose haplotype estimation was ambiguous (i.e. heterozygous at both SNPs). Haplotype [A-T] carriers (red) have surprisingly small (i.e., 20-22) KIV number on the predominantly expressed isoform.

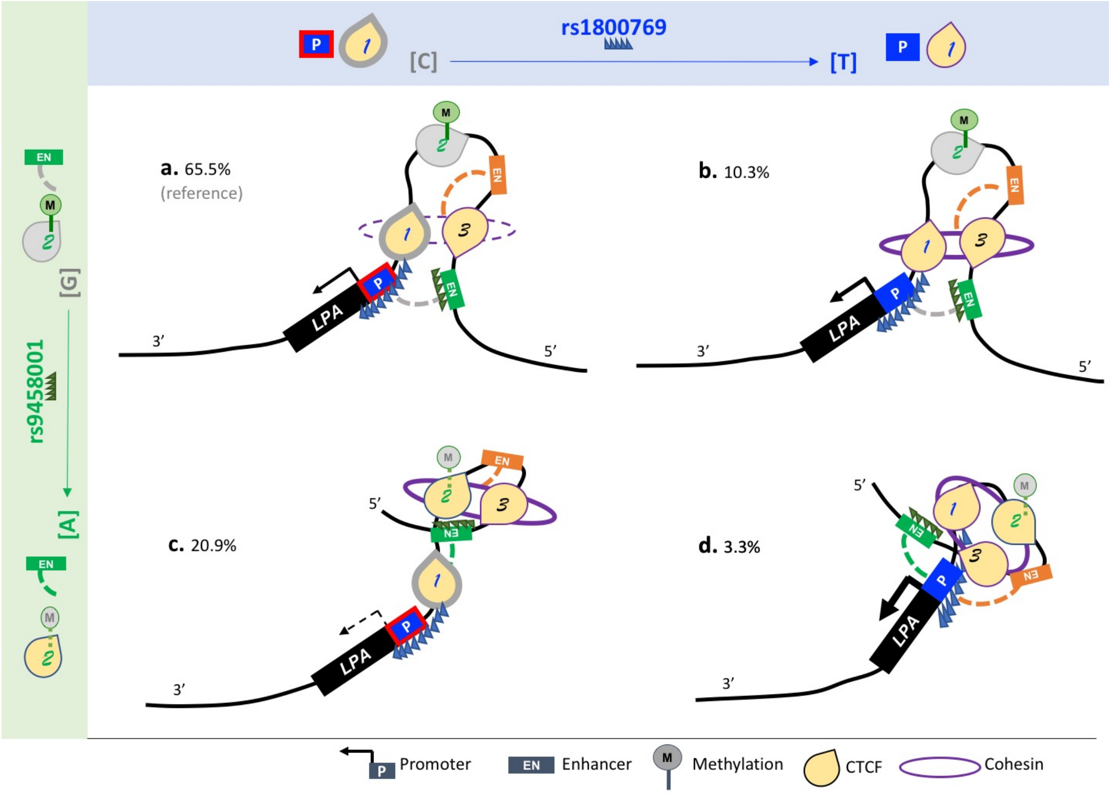


**Supplementary Figure 10**

**Hypothetical co-regulation of LPA gene expression by rs1800769 and rs9458001.**

We propose that the summarized expression activity (thickness of the black arrow on the promoter) of the gene is determined both by the innate promoter activity and the additional expression activity owing to the active enhancer-promoter binding. The four panels **a-d** display the four scenarios of promoter-enhancer interactions for Lp(a) transcribed on the reverse strand in the direction 5' to 3' for respective haplotypes composed of rs1800769-rs9458001 [C-G], [T-G], [C-A] and [T-A]. rs1800769 is located at the promoter region of *LPA* (rectangles labelled with 'P' and filled in blue) as well as with several SNPs in high LD of it located inside the gene body (blue saw shapes); rs9458001 and SNPs in high LD of it (green saw shapes) are located at the intergenic region upstream of *LPA* promoter (green). The three CTCF-DNA binding sites indicated by the UCSC genome browser (Supplementary Note V) are displayed as yellow drops, with rs1800769 LD region overlapped at position 1 (label text in blue) and rs9458001-mediated methylation overlapped at position 2 (label text in green). The two massive signals for the enhancer elements indicated from UCSC genome browser (Supplementary Note V) are displayed as rectangles labelled with 'EN', with specifically the one at the rs9458001 local region colored in green. Both SNPs are bi-functional. Rs1800769[T] compared to [C] allele has stronger promoter activity[3] (non-red outline), and meanwhile has higher local CTCF-DNA binding affinity at position 1 (non-grey outline) (ENCODE[4], Supplementary Table 10). Rs9458001[A] compared to [G] leads to stronger co-activators recruiting at the green enhancer (hypothesized based on enhancer histone signal, Supplementary Table 10, HaploRegv4[5]), and meanwhile increase of CTCF-DNA binding at position 2 (via decrease local methylation[6], Methods, Supplementary Table 11). Cohesin and CTCF differentially affect the topological architecture of DNA loops[7-9]. Panel **a** (65.5%) reflects the most frequent haplotype and is the reference scenario, with weak promoter activated with the standard effect of green enhancer (grey dotted line), and unstable cohesin binding with CTCF 1 and 3 (possible insulation of orange enhancer). Compared to the reference (**a**), in senario **b** (10.3%), the promoter activity is higher due to the change of rs1800769[T] allele, but binding of cohesion-CTCFs is more stable, which results in only slightly elevated levels of expression. Compared to the reference (**a**), in scenario **c** (20.9%), the CTCF-DNA binding at position 2 is activated due to the change of rs9458001[A] allele. As the CTCF-binding at position 1 is still weak, the cohesin preferably combines together the CTCF 2 and 3. The orange enhancer is securely blocked. The green enhancer near position 3, although more activated, is ineffective at interacting with the promoter because of being tightly bound to the cohesin and therefore is far away from the promoter; additionally it is being insulated by CTCF 1 (although unstable) from interaction. This leads to dramatically reduced expression. Lastly, in panel **d** (3.3%), not only is the promoter activity higher, but also both enhancers are exposed (cohesin bring all three stably bound CTCFs together) to the promoter element and thus finally leads to exceptionally high expression activity.

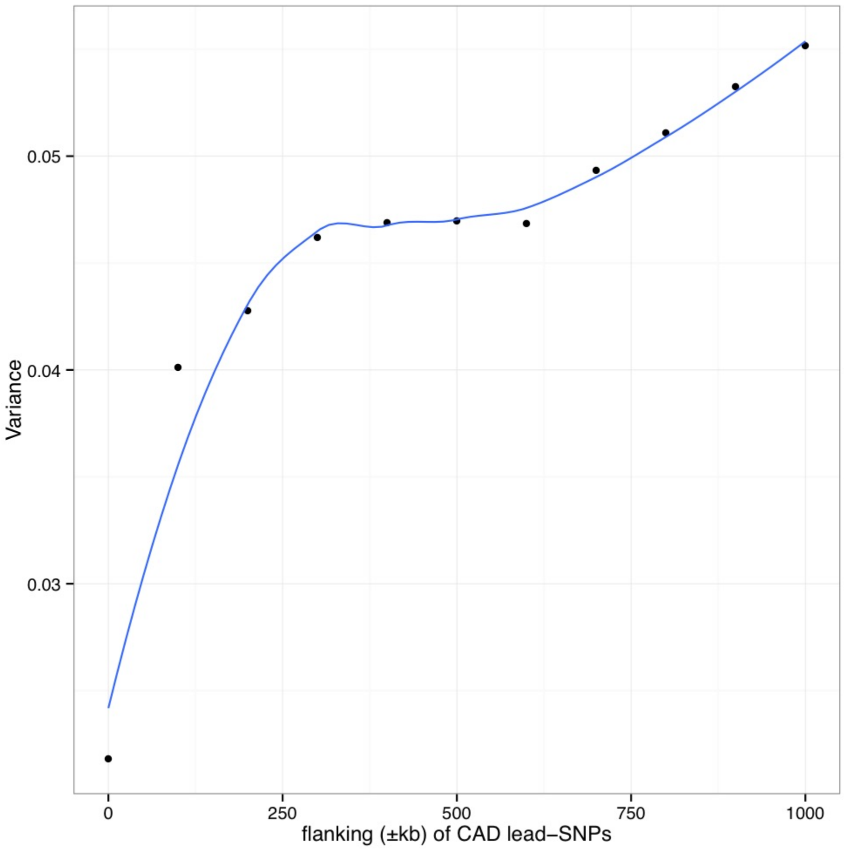


**Supplementary Figure 11**

**Increased variance explained by physical expansion around the known CAD lead SNPs reported from GWAS studies.**

All variants in the flanking region around the lead SNPs at 56 loci with available genotypes in nine CAD case-controls studies were extracted. LDAK tool was used to calculate the LD-adjusted kinship matrix among all individuals, which was then forwarded to GCTA to estimate the SNP-based heritability of CAD in the measurement of the total variance explained in liability model (assuming a prevalence of CAD as 5%). The x-axis represents the flanking range of known loci, from ±100kb to ±1mb progressively with steps of 100kb. The y-axis represents the variance of CAD risk that could be explained by the given SNPs in the liability model. Indeed, variance explained including the flanking regions achieved 0.47 around the step at ±500kb, while variance explained of the lead SNPs only (±0kb) was 0.22, which was only a 46.5% proportion (0.22/0.47).

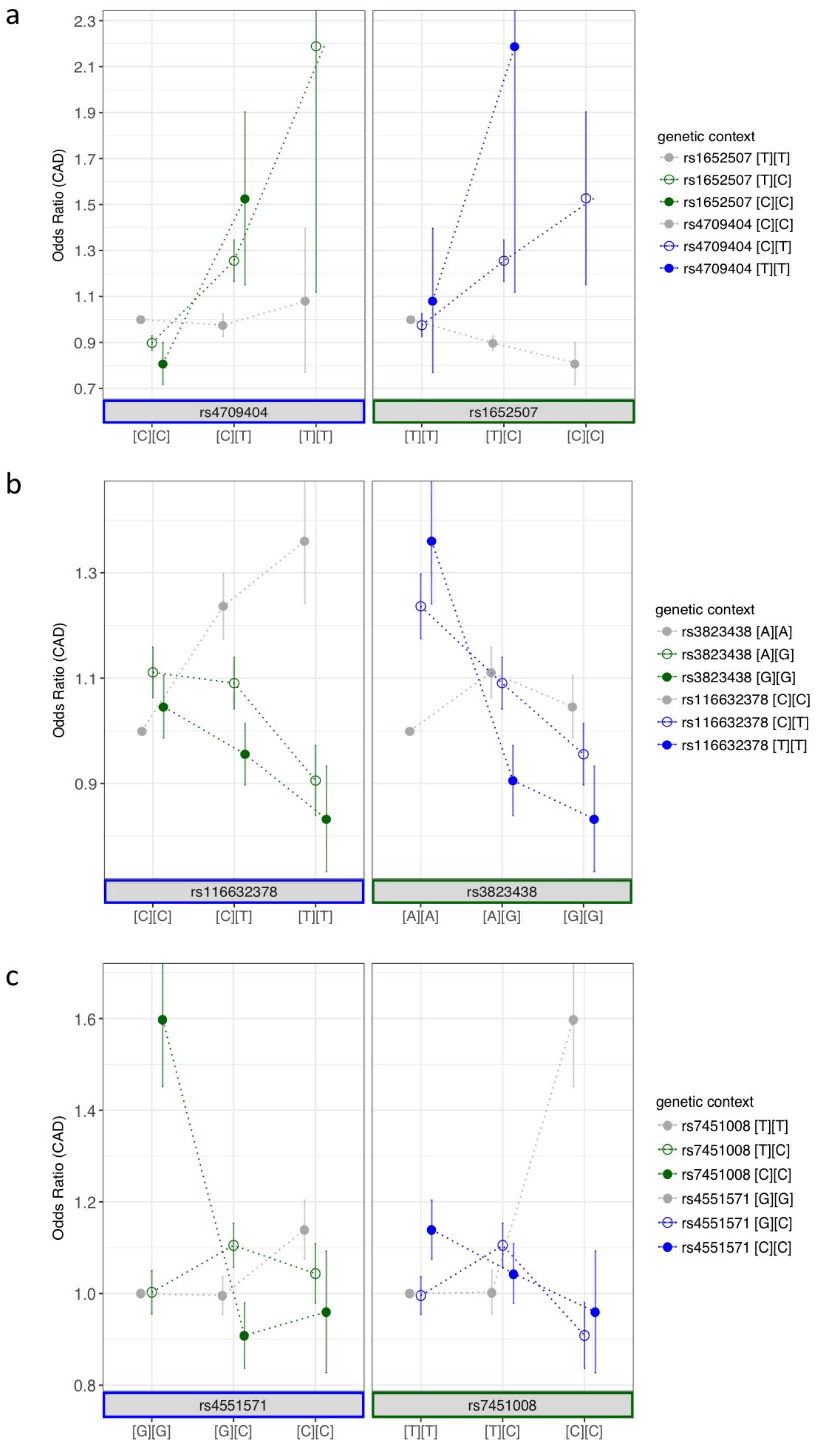


**Supplementary Figure 12**

**Relative effect sizes for subgroups of individuals with different genotype context given the other 3 candidate SNP pairs.**

Odds ratio of CAD and standard errors are displayed in biallelic perspective for 9 genotype combinations, with the reference genotype combination set as the most frequent genotype. **a.** cis SNP-pair rs4709404 and rs1652507 in [T] dosage and [C] dosage model. **b.** cis SNP-pair rs116632378 and rs3823438 in [T] dosage and [G] dominant model. **c**. trans SNP-pair rs4551571 and rs7451008 in [C] dominant and [C] recessive model. Epistasis is demonstrated as genetic effects of one SNP depending on the genetic context of the other SNP.

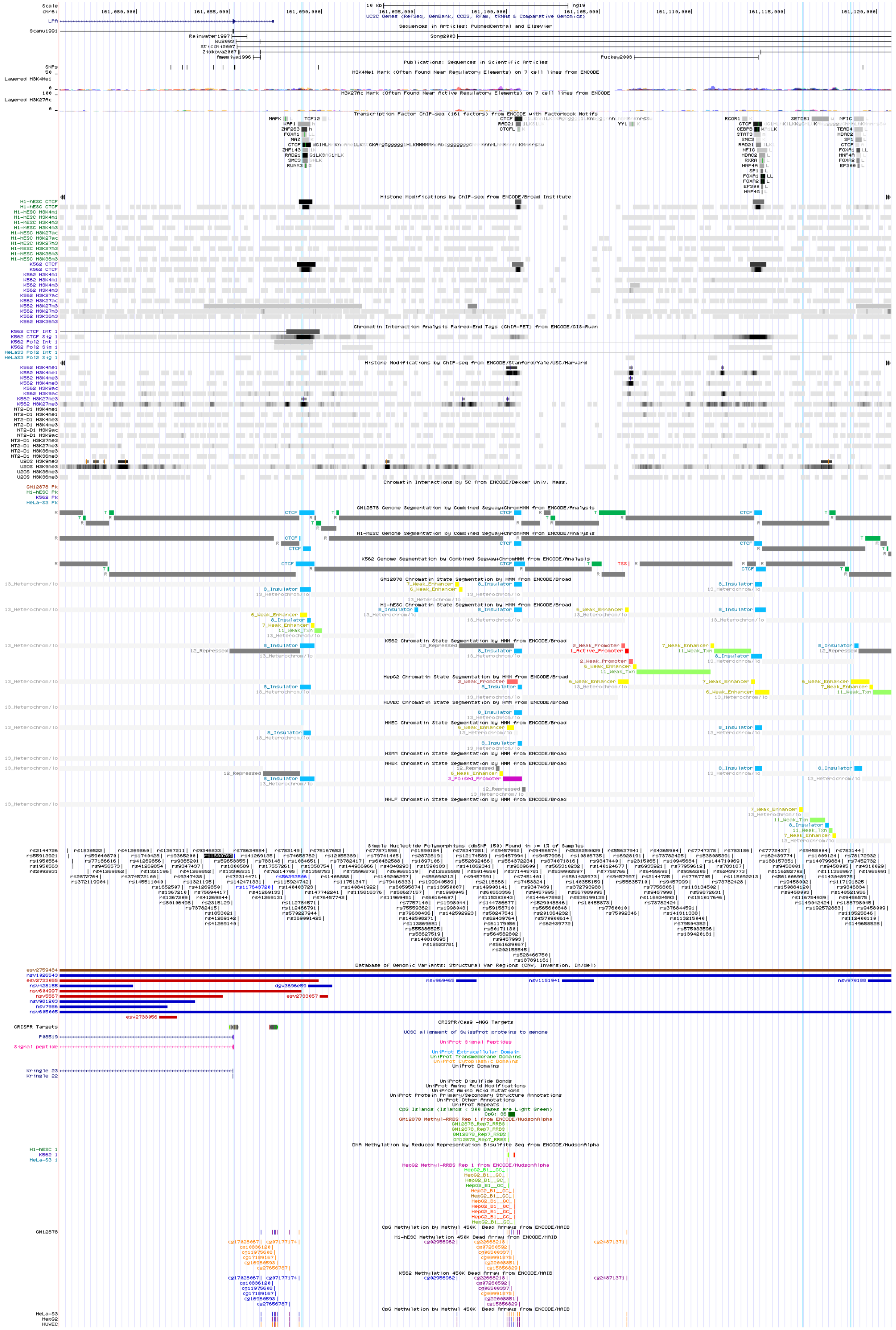
Scale
chr6:
10 kb
hg19
UCSC Genes (RefSeq, GenBank, CCDS, Rfam, tRNAs & Comparative Genomics)
LPA
Sequences in Articles: PubmedCentral and Elsevier
Publications: Sequences in Scientific Articles
H3K4Me1 Mark (Often Found Near Regulatory Elements) on 7 cell lines from ENCODE
H3K27Ac Mark (Often Found Near Active Regulatory Elements) on 7 cell lines from ENCODE
Transcription Factor ChIP-seq (161 factors) from ENCODE with Factorbook Motifs
Histone Modifications by ChIP-seq from ENCODE/Broad Institute
Chromatin Interaction Analysis Paired-End Tags (ChIA-PET) from ENCODE/GIS-Ruan
Histone Modifications by ChIP-seq from ENCODE/Stanford/Yale/USC/Harvard
Chromatin Interactions by 5C from ENCODE/Dekker Univ. Mass.
GM12878 Genome Segmentation by Combined Segway+ChromHMM from ENCODE/Analysis
H1-hESC Genome Segmentation by Combined Segway+ChromHMM from ENCODE/Analysis
K562 Genome Segmentation by Combined Segway+ChromHMM from ENCODE/Analysis
GM12878 Chromatin State Segmentation by HMM from ENCODE/Broad
H1-hESC Chromatin State Segmentation by HMM from ENCODE/Broad
K562 Chromatin State Segmentation by HMM from ENCODE/Broad
HepG2 Chromatin State Segmentation by HMM from ENCODE/Broad
HUVEC Chromatin State Segmentation by HMM from ENCODE/Broad
HMEC Chromatin State Segmentation by HMM from ENCODE/Broad
HSMM Chromatin State Segmentation by HMM from ENCODE/Broad
NHEK Chromatin State Segmentation by HMM from ENCODE/Broad
NHLF Chromatin State Segmentation by HMM from ENCODE/Broad
Simple Nucleotide Polymorphisms (dbSNP 150) Found in >= 1% of Samples
Database of Genomic Variants: Structural Var Regions (CNV, Inversion, In/del)
CRISPR/Cas9 -NGG Targets
UCSC alignment of SwissProt proteins to genome
UniProt Signal Peptides
UniProt Extracellular Domain
UniProt Transmembrane Domains
UniProt Cytoplasmic Domains
UniProt Domains
UniProt Disulfide Bonds
UniProt Amino Acid Modifications
UniProt Amino Acid Mutations
UniProt Protein Primary/Secondary Structure Annotations
UniProt Other Annotations
UniProt Repeats
CpG Islands (Islands < 300 Bases are Light Green)
GM12878 Methyl-RRBS Rep 1 from ENCODE/HudsonAlpha
DNA Methylation by Reduced Representation Bisulfite Seq from ENCODE/HudsonAlpha
HepG2 Methyl-RRBS Rep 1 from ENCODE/HudsonAlpha
CpG Methylation by Methyl 450K Bead Arrays from ENCODE/HAIB
H1-hESC Methylation 450K Bead Array from ENCODE/HAIB
K562 Methylation 450K Bead Array from ENCODE/HAIB
CpG Methylation by Methyl 450K Bead Arrays from ENCODE/HAIB

**Supplementary Figure 13**

**Overview of functional annotations for the genomic region in the vicinity of rs1800769 and rs9458001.**

Interactively selected annotations for the spanning genomic region (hg19, chr6:161,050,000-161,120,000) downloaded from UCSC genome browser (https://genome.ucsc.edu). The region in between the first two blue vertical lines corresponds to the positions where rs1800769 and its proxies are located. The region in between the third and fourth blue vertical lines corresponds to the positions where rs9458001 and its proxies are located.

### Supplementary Notes


**I. The broad-sense CAD susceptibility regions**

**II. Genotype processing for 10 CAD case-control studies**

**III. Clinical significance of epistasis**

**IV. Size of Apo(a) isoforms and KIV-2 CNV numbers**

**V. Hypothesized mechanism of epistasis**

**VI. Evidence against haplotype effect**


#### I. The broad-sense CAD susceptibility regions

We focused our analysis at loci with previous evidence of genome-wide association with CAD in order to restrict the number of variants for testing of statistical epistasis with the aim to enhance speed and the chance of positive finding. Firstly, we defined regions of CAD susceptibility in a broad-sense, i.e., flanking regions of known CAD risk loci, given that multiple independent signals at the surrounding region of known CAD lead-SNPs could explain additional heritability to CAD[10,11]. To achieve this, we collected the lead SNPs from the 56 published CAD susceptibility loci[10,12], and calculated the SNP-based heritability for all SNPs within a certain flanking range of known loci, from ±100kb to ±1mb progressively with steps of 100kb.

For heritability calculation, all variants in the flanking region around the lead SNPs at 56 loci with available genotypes in nine CAD case-controls studies were extracted (GerMIFSVI not included at this time). Imputed genotypes or proxy variants were used where necessary. The LDAK tool[13] was used to calculate the LD-adjusted kinship matrix among all individuals, which was then forwarded to software GCTA[14] to estimate the SNP-based heritability of CAD in the measurement of the total variance explained in liability model (assuming a prevalence of CAD as 5%). As expected, an incremental increase in the heritability was observed with the enlargement of the flanking region (Supplementary Fig 11). The increase of heritability explained by the regions increased profoundly from ±100kb until ±500kb but largely attenuated afterwards. Therefore, we decided ±500kb as a balanced threshold for the flanking range surrounding the known loci (i.e., the broad-sense CAD susceptibility regions), which could meanwhile maximize the heritability covered by the regions and minimize the computational burden.

### II. Genotype processing for 10 CAD case-control studies

| Study | Array Platform |
|---|---|
| GerMIFSI | Affymetrix Mapping 500K Array Set |
| GerMIFSII | Affymetrix Genome-Wide Human SNP Array 6.0 |
| GerMIFSIII | Affymetrix Genome-Wide Human SNP Array 5.0/6.0 |
| GerMIFSIV | Affymetrix Genome-Wide Human SNP Array 6.0 |
| GerMIFSV | Illumina HumanOmniExpress/Omniuni_2.5 |
| LURIC | Affymetrix Genome-Wide Human SNP Array 6.0 |
| WTCCC | Affymetrix Genome-Wide Human SNP Array 6.0 |
| MIGEN | Affymetrix Mapping 500K Array Set |
| Cardiogenics | Illumina Human660W-Quad |
| GerMIFSVI | Illumina PsychChip_v1-1 |

*Supplementary Table 1. Gentoype array platforms of ten CAD case-control studies used in the discovery stage.*

The 10 case-control studies of coronary disease were originally genotyped with the corresponding arrays (Supplementary Table 1). The following pre-imputation quality control (QC) criteria were taken: individual call rate >= 0.98, SNP call rate > 0.98, minor allele frequency (MAF) > 0.01, concordant recorded and genotype-derived gender, population outliers excluded (deviate beyond mean ± 5×(standard deviation (SD)) for top two dimensions from the multidimensional scaling (MDS) analysis, PI_HAT < 0.0625 (individuals more distant away than fourth-degree relatives) in the identity-by-descent (IBD) analysis, heterozygosity rate within mean ± 3×SD, and deviation from Hardy-Weinberg Equilibrium (HWE) p > 1e-6.

After genotype QC, we used all individuals from the 1000 Genomes Phase 1 Version 3 reference panel (1000G) to impute the genotypes. Haplotypes were firstly pre-phased from genotypes with SHAPEIT2 haplotype estimation tool to generate the best guess haplotypes based on the given genotypes. Then the best guess haplotypes were forwarded to IMPUTE2 for imputation. Finally, the following post-imputation QC criteria were taken: SNP call rate > 0.98, MAF > 0.05, Hardy-Weinberg p > 1e-5, INFO score >= 0.8.

In the process of our work, the HRC reference dataset[15] was released as the largest available haplotype reference panel for imputation of variants in populations of European ancestry. The coverage of SNPs in our data based on HRC imputation were higher compared to 1000G, thus we repeated the same analysis based the HRC v1.6 imputation, with the hope of fine-mapping the lead-SNP-pair. The same pre-imputation and post-imputation QC criteria were applied as for 1000G. The imputation procedures were conducted through Sanger Imputation Server (https://www.sanger.ac.uk/science/tools/sanger-imputation-service).

#### III. Clinical significance of epistasis

Paradoxical context-dependent effect sizes were observed for all statistical interacting SNP pairs, in their genetic model for effect alleles (Fig 3, Supplementary Fig 12). In other words, it seems the usually tested effect size in the conventional GWAS analysis (Supplementary Table 2) only arbitrarily assumed the genetic model (i.e., additive), and represented an averaged effect size against various genetic backgrounds but masked the actual largely paradoxical effects dependent on the different genetic context of each other.

In this study, we prioritized our four candidate SNP- pairs that hit all statistical criteria for candidate epistasis, and finally focused on the lead SNP-pair rs1800769-rs9458001 in dosage-dosage model for investigation of biological epistasis. In this model, the effects of two SNPs on CAD susceptibility are truly unveiled only when dissected against the mutual genetic context of each other. Univariately – against all other genetic backgrounds – rs9458001[A] and rs1800769[T] had on average neutral effects, but studying their interaction unmasked the largely divergent and opposite effects dependent on the different genetic context of each other.

Irrespectively of the partially unknown mechanism, this epistasis pair has extraordinary clinical significance. A single haplotype [T-A] which compose ~3% in the population, displayed extremely high risk of CAD ($P=5.6 \times 10^{-11}$, OR=1.84, Fig 2b, Supplementary Table 6), even much higher than the so-far reported genome-wide significant CAD lead SNP rs10455872 at the same locus (OR 1.31)[11] (Supplementary Fig 7). The highest risk subgroup in the ten case-controls studies was rs1800769[TT] - rs9458001[AA] carriers (n=7), which in fact, were exclusively CAD cases. Carriers of the [T-A] haplotype not only exhibits far higher promoter activity (Fig 3b), but also a predominance of smaller KIV-2 CNV size (i.e. 20-22) (Supplementary Fig 9). Compared to the reference haplotype [C-G] (21.3 mg/dl), haplotype [T-A] had 49.6 mg/dl higher Lp(a) level (Supplementary Table 12,13). The only [T/T] - [A/A] genotype carrier in the LURIC study, had almost 10-fold higher Lp(a) levels (224 mg/dl) compared to the KORA population mean of 22.05 (Supplementary Table 12). Further efforts of incorporating epistasis information would be worthwhile to enhance the development of more accurate models for genetic risk prediction, and potential possibilities of genetic therapy.

**IV. Size of Apo(a) isoforms and KIV-2 CNV numbers**

Apo(a) isoforms were determined by sodium dodecyl sulfate-agarose gel electrophoresis (SDS agarose) under reducing conditions as described in Kronenberg et al[16]. Electrophoresis was followed by immunoblotting using the monoclonal antibody 1A2 for detection of apo(a) isoforms. Given the large number of alleles present in the population, >90% of the individuals in a population are heterozygous on DNA level. However, only about 60-70% present both isoforms in plasma. Larger isoforms tend to be non-expressed due to overly long residence in the endoplasmatic reticulum. Short isoforms are produced more efficiently per time unit and thus expressed at a higher level, commonly contributing to a higher extent to the circulating Lp(a) and thus present a stronger band in the Western blot (>50% of the total intensities of both band). Given the limit of resolution of about +/- 1 KIV repeat, presenting only one band may be either truly homozygous, "rather homozygous" (e.g., situation A: When two people have KIV-2 repeat numbers 20 and 21, the second isoform may not be expressed at all. In this study, all statistical models use the predominantly expressed isoform for isoform-based adjustment.

**V. Hypothesized mechanism of epistasis**

Towards the potential molecular mechanism of rs1800769-rs9458001 epistasis on *LPA* gene regulation, besides multiple functional annotations for the two SNPs from literature, Annovar[4], ENCODE[4], HaploReg[5], etc. (see Methods), we also resorted to the UCSC genome browser (https://genome.ucsc.edu) and interactively selected annotations for the spanning genomic region (hg19, chr6:161,075,834-161,120,798, saved as Supplementary Fig 13). *LPA* gene is transcribed on the reverse strand in the direction 5' to 3'. rs1800769 and SNPs in high LD of it are located near the promoter region of *LPA* as well as with several SNPs in high LD of it located inside the gene body (between the first and second vertical lines); rs9458001 and SNPs in high LD of it are located at the intergenic region upstream of *LPA* promoter and near the enhancer elements (between the third and fourth vertical lines). The three CTCF-DNA binding sites indicated from UCSC genome browser are located at position 1, with rs1800769 LD region overlapped; at position 2, with rs9458001-mediated CpG methylation (chr6:161100092-161100456, ~15kb upstream of rs1800769 and downstream of rs9458001, as based on two independent methylation array analyses[6], Methods Supplementary Table 11) overlapped; and at position 3, just before the rs9458001 LD region.

We hypothesize that the summarized expression activity of the *LPA* gene is affected by the genetically mutually dependent coordination of two SNPs, and is determined both by the innate promoter activity and the additional activity owing to active enhancer-promoter interaction (Supplementary Fig 10). Both SNPs are bi-functional. Rs1800769[T] compared to [C] allele has stronger promoter activity[3] , and meanwhile has higher local CTCF-DNA binding affinity at position 1 (ENCODE[4], Supplementary Table 10). Rs9458001[A] compared to [G] leads to stronger co-activators recruiting at the green enhancer (hypothesized based on enhancer histone signal, Supplementary Table 10, HaploRegv4[5]), and meanwhile increase of CTCF-DNA binding at position 2 (via reducing methylation of the local CpG island, which in turn affects the chance of barrier-free CTCF-DNA binding). Furthermore, it is well-known that mammalian genomes are folded at multiple scales and each scale highlights an important interplay between structure and function, in which CTCF and cohesin are critical regulators[7-9]. We hypothesize that their allele-specific effects on CTCFs are intermediary to the regional CTCF-cohesin topological structure[8], and thereby regulate the enhancer-promoter interaction and gene expression in a context-dependent way (Supplementary Fig 10, a-d). While admitting that the precise molecular mechanisms need further substantiation by functional studies, our proposed mechanism for this epistasis pair appears to be plausibly supporting that both SNPs should display pleiotropic effects so as to compose together a prerequisite for epistasis[17].

In a broader perspective, epistasis has been suggested to play a prominent role in modulating evolution[18-21]. One involvement is co-adaption (which originally refers to that two (or more) genes undergo adaptation as a pair (or group)), where positively interacting (co-adapted) alleles tend to be co-inherited[22] under favorite selection, thereby making alleles with higher Darwinian fitness more common over time. Genomic imprinting has been suggested as a way to choose which of the two alleles at a locus is preferentially expressed and to coordinate the variants that interact epistatically[23]. Coincidentally and interestingly, there is a special phenomenon in apo(a) expression in that often one allele is predominantly expressed (Supplementary Note IV). Indeed, in our case, the most frequent haplotype rs1800769[C]-rs9458001[G] comes with low CAD risk.

Besides co-adaptation, epistasis has also been suggested to facilitate genetic robustness (the insensitivity of organisms to the impact of mutations)[17]. In our case, the Lp(a) concentration is notoriously known for its highly right skewed distribution in the population[24], where the majority of individuals display rather low levels of Lp(a), which thus seems to be under strict control. However, a small proportion in the skewed distribution curve display especially high Lp(a) levels. Undoubtedly, further investigation is needed to corroborate as to whether imprinting and co-adaptation is involved mechanistically, or whether the exceptionally high Lp(a) levels is due to the escape from the robust regulation.

**VI. Evidence against haplotype effect**

To further corroborate the concept that the high CAD risk of the haplotype rs1800769[T]-rs9458001[A] (or the genotype combination rs1800769[T/T]-rs9458001[A/A]) is due to the result of their epistasis effect, rather than the additive effect of a potentially masked haplotype, we scrutinized all the genotyped SNPs in between these two SNPs specifically for all individuals in this high risk group (i.e., with the exact rs1800769[T/T] and rs9458001[A/A] genotype combination), and additional LD structure information was characterized (Supplementary Table 14). It was observed that rs1800769 (proxies) and rs9458001 (proxies) are located in distinct haplotype blocks, and that SNPs in-between have inconstant allele frequencies and heterozygous genotypes across these high risk individuals, spanning across two different haplotype blocks involving three recombination breakpoints (Supplementary Table 14). These provided further evidence arguing against the possibility of signal derived from an untagged rare variant instead of the two SNPs.

In addition, we also tested directly whether a putative haplotyped tagged by a haplotype derived rs1800769 (proxies) and rs9458001 (proxies) provided a better model fit as compared to the epistasis model found in this study. To this end, we were able to use six of the ten original (GerMIFSII, GerMIFSV, LURIC, WTCCC, Cardiogenics and GerMIFSVI). In these studies, we had genotyped data available for the rs1800769 (proxies) and rs9458001 (proxies). We then compared model performance for a logistic model with haplotypes estimated using the R package *haplo.stats* (https://CRAN.R-project.org/package=haplo.stats), and found a model allowing for the additional inclusion of the interaction term to be a significantly better fit to the data ($P=7.67\times10^{-6}$) even after forcing the haplotypic effects to be included in the model (Supplementary Table 15). Using a stepwise backward AIC based algorithm none of the estimated haplotypic effects were retained in the final model, with the interaction effect clearly remaining significant ($P= 2.081\times10^{-09}$) for the six studies combined. Even allowing for interactions between the estimated haplotypes did not alter the results of the stepwise (Supplementary Table 15). For these analyses, we used gender, study and 10 MDS components as covariates. We take these results and considerations jointly as evidence that phased information is unlikely the cause of the results seen. Thus, indirectly, the effect of a haplotype or a single rare variant tagged by it is improbable.
